# Chimeric GPCRs mimic distinct signaling pathways and modulate microglia responses

**DOI:** 10.1101/2021.06.21.449162

**Authors:** Rouven Schulz, Medina Korkut-Demirbaş, Gloria Colombo, Sandra Siegert

**Affiliations:** Institute of Science and Technology (IST) Austria, Am Campus 1, 3400 Klosterneuburg, Austria

**Keywords:** G protein coupled receptor (GPCR), DREADD, β2-adrenergic receptor (β2AR/ADRB2), microglia, inflammation, GPR65, GPR109A/HCAR2

## Abstract

G protein-coupled receptors (GPCRs) regulate multiple processes ranging from cell growth and immune responses to neuronal signal transmission. However, ligands for many GPCRs remain unknown, suffer from off-target effects or have poor bioavailability. Additional challenges exist to dissect cell type-specific responses when the same GPCR is expressed on different cells within the body. Here, we overcome these limitations by engineering DREADD-based GPCR chimeras that selectively bind their agonist clozapine-N-oxide (CNO) and mimic a GPCR-of-interest. We show that the chimeric DREADD-β2-adrenergic receptor (β2AR/ADRB2) triggers comparable responses to levalbuterol on second messenger and kinase activity, post-translational modifications, and protein-protein interactions. Moreover, we successfully recapitulate β2AR-mediated filopodia formation in microglia, a β2AR-expressing immune cell that can drive inflammation in the nervous system. To further dissect microglial inflammation, we compared DREADD-β2AR with two additionally designed DREADD-based chimeras mimicking GPR65 and GPR109A/HCAR2, both enriched in microglia. DREADD-β2AR and DREADD-GPR65 modulate the inflammatory response with a similar profile as endogenously expressed β2AR, while DREADD-GPR109A had no impact. Our DREADD-based approach allows investigation of cell type-dependent signaling pathways and function without known endogenous ligands.

## Introduction

The translation of extracellular signals into an intracellular response is critical for proper tissue function. G protein-coupled receptors (GPCRs) are key mediators in this process with their strategic placement at the cell membrane to bind diverse molecule classes ^1,2^. Successful ligand-GPCR interaction triggers intracellular signaling cascades with far-reaching impacts on cell functions like growth, migration, metabolism, and cell-cell communication ^3,4^. Approximately 35% of all *food and drug administration* (FDA)-approved drugs target GPCR ^5,6^, stressing their importance for biomedical research and drug development. However, major challenges exist in investigating GPCR signaling. First, GPCRs often have unidentified ligands, including more than 100 potential drug targets and the majority of olfactory receptors ^5,7,8^. Second, GPCR expression and signaling are cell type-specific. For example, β2-adrenergic receptor (β2AR/ADRB2) modulates inflammation in immune cells ^9^, relaxes smooth muscle in bronchial tubes ^10^, and impacts pancreatic insulin secretion and hepatic glucose metabolism ^11^. Such response diversities hinder dissecting cell type-dependent effects *in-vivo*. Third, GPCR ligands often suffer from poor bioavailability or cause off-target effects. For instance, norepinephrine acts as ligand for β2AR but can also activate other adrenoceptors in the central nervous system ^12^. Therefore, novel strategies are required to overcome the limitations of unknown or unsuitable ligands and simultaneously allow selective investigation of GPCR signaling in a cell type-of-interest.

So far, over 800 GPCRs are known, which are structurally conserved with seven transmembrane helices (TM) connected by three extracellular (ECL) and intracellular (ICL) loops ^13^. Ligand binding involves N-terminus, ECLs, and TM domains and consequently triggers ICL interaction with heterotrimeric G proteins. These G proteins are composed of α- and βγ-subunits that act as effectors on downstream signaling partners ^13^. Specific subunit recruitment of either Gα_s_, Gα_q_ or Gα_i_ activates defined canonical pathways ^14^. Besides ICLs as critical components for proper GPCR signal transduction ^15–17^, the C-terminus interacts with β-arrestins, which contribute to receptor desensitization ^18–20^ and kinase recruitment ^21–25^. Several studies have exploited the concept of ligand binding and signaling domains to control GPCR function ^26–31^. Airan *et al.* generated light-inducible GPCR chimeras that mimic the signaling cascades of distinct GPCRs ^27^. However, caveats exist with this optogenetic approach, as it relies on strong light stimulation which induces phototoxicity ^32–35^. Additionally, light exposure *in-vivo* requires invasive procedures that will disrupt tissue integrity and alter the response of resident immune cells ^36^. Yet, immune cells are interesting targets for studying GPCR signaling as their function and ability to induce inflammation is tightly controlled by these receptors ^37–39^. Several immune cells such as circulating leukocytes and lymphocytes are not confined to any light-accessible tissue and therefore cannot be manipulated through light-inducible GPCRs.

Here, we designed chemical-inducible GPCR chimeras based on the DREADD system (Designer Receptor Exclusively Activated by Designer Drugs) ^40–42^. DREADDs are modified muscarinic acetylcholine receptors, which are inert to their endogenous ligand acetylcholine and respond to clozapine-N-oxide (CNO), a small injectable compound with minimal off-target effects and suitable bioavailability for *in-vivo* usage ^43^. We identified the ligand binding and signaling regions of DREADD and 292 GPCRs. This enabled us to engineer CNO-responsive chimeras with β2AR being our proof-of-concept candidate due to its well-known ligand and broad physiological importance ^10,11^, which includes modulating inflammation in various immune cells ^9^ such as microglia ^44^. After exchanging the corresponding signaling domains, we established that DREADD-β2AR fully recapitulated the signaling pathways of levalbuterol-stimulated non-chimeric β2AR including the impact on microglia motility ^45^. Finally, we identified immunomodulatory effects of DREADD-β2AR and two additionally established DREADD chimeras for the microglia-enriched GPR65 and GPR109A/HCAR2. This underlines that our approach can be applied to different GPCRs-of-interest allowing cell type-targeted manipulation of GPCR signaling.

## Results

### Establishing a library for *in-silico* design of DREADD-based GPCR chimeras

Microglia are tissue-resident macrophages of the central nervous system. They maintain homeostasis during physiological conditions and induce an inflammatory response upon tissue damage and pathogen encounter ^46,47^. GPCRs are critical for these functions as they allow fast adaption to local perturbations. To identify which GPCRs are selectively enriched in microglia, we compared GPCR expression across different cell types in a previously established retina transcriptome database ^48^. We found approximately one-third of the most abundant GPCRs enriched in microglia, which also included the well-defined β2-adrenergic receptor (β2AR) ^9–11,44,45,49^, making it a prime candidate for establishing our strategy (**Fig.1a**).

To design CNO-responsive DREADD-based chimeras mimicking a GPCR-of-interest (**Fig.1b**), we first identified GPCR ligand binding and signaling domains including either N-terminus, extracellular loops (ECLs) and transmembrane helices (TMs), or intracellular loops (ICLs) and C-terminus, respectively (**Fig.1c**). We performed multiple protein sequence alignment using the established domains of bovine rhodopsin (RHO) ^27^ as reference. We aligned rhodopsin with the CHRM3-based DREADD (from now on referred to as DREADD), human β2-adrenergic receptor (β2AR), and 292 other potential GPCRs-of-interest (**Fig.1d**). As internal controls, we included human α1-adrenergic receptor (hα1AR) and hamster β2-adrenergic receptor (hamβ2AR) and confirmed that our alignment successfully reproduced the rhodopsin-based chimeras from *Airan et al.* ^27^. To further verify alignment accuracy, we utilized the TMHMM algorithm, which predicts TM domains in a protein sequence ^50^. For all GPCRs shown in **Figure 1d** and **Supplementary Figure S1**, predicted TMs displayed the expected tight flanking of ICLs and C-termini identified by our alignment. The only exception was GPR109A, where TMHMM failed to identify the seventh TM with sufficient certainty. Occasionally, minor deviations from seamless flanking occurred but were within the observed range for the reference and internal controls (**Supplementary Fig.S1**). Finally, we exploited published crystal structures for three key GPCRs: the alignment reference RHO; the DREADD, for which we used rat CHRM3 as surrogate with over 90% sequence similarity; and our primary GPCR-of-interest β2AR. We mapped our identified ligand binding and signaling domains together with predicted TMs on these crystal structures and found that they closely matched the expected extracellular, transmembrane or intracellular locations (**Fig.1e**). These results suggest that our alignment correctly predicts GPCR domains and serves as a library for generating DREADD-based GPCR chimeras.

**Figure 1:**
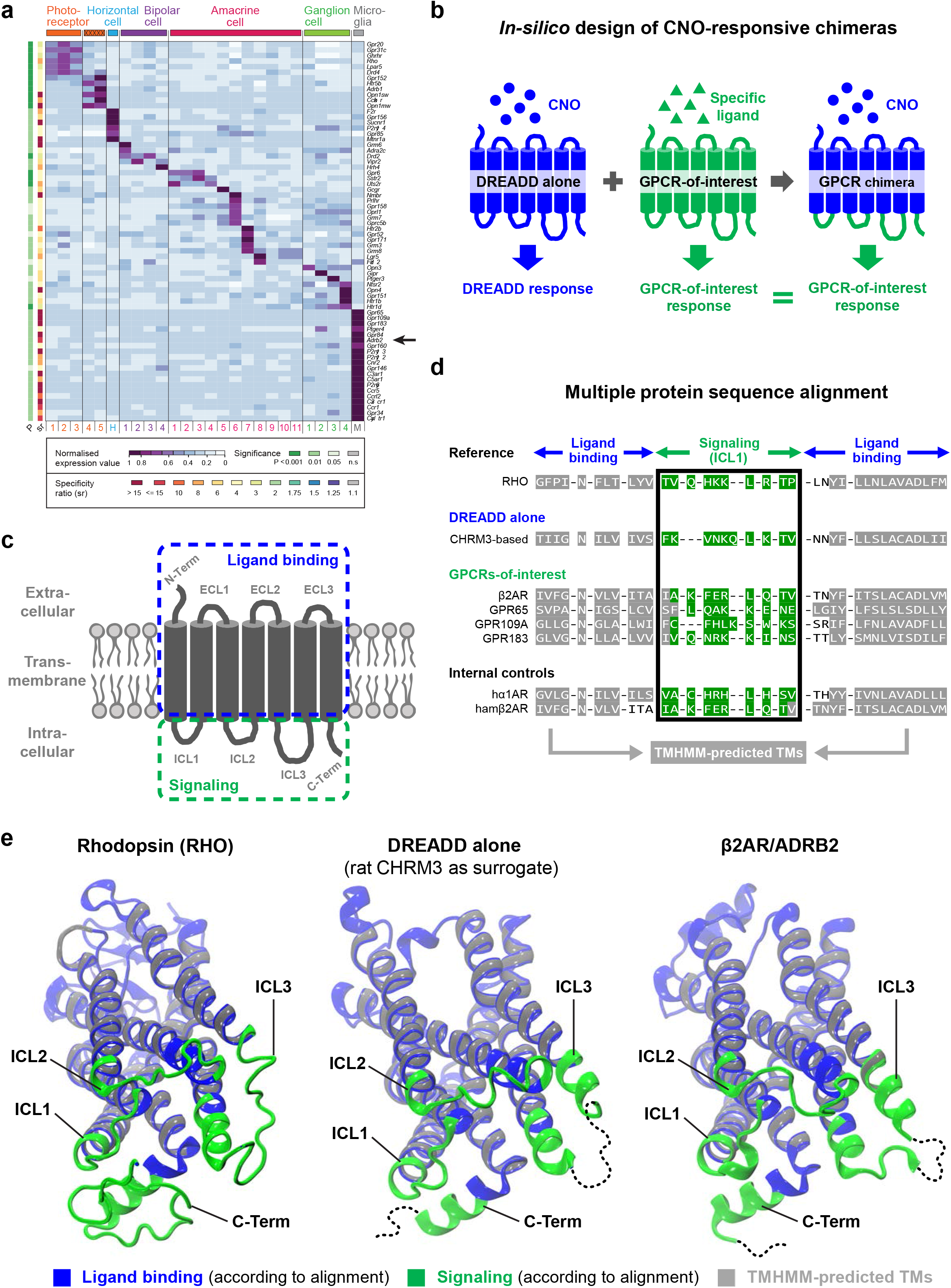
GPCR domains can be identified *in-silico* to engineer chimeric receptors. **a:** GPCR gene expression analysis across different cell types in the mouse retina. Columns represent distinct cell types and show clusters of selectively enriched GPCRs. Purple indicates high gene expression. *P*-value and specificity ratio (s.r.) are color-coded as indicated in the figure and described in Siegert *et al.* ^48^. Arrow points to β2AR/ADRB2. **b**-**c:** Schematic of GPCR domains and chimera design. **b:** The intracellular domains of a DREADD (blue) are replaced with the intracellular domain of a GPCR-of-interest (green), generating a chimeric receptor that induces the signaling cascade of the GPCR-of-interest (green) upon CNO stimulation. **c:** GPCRs consist of seven transmembrane domains, three extracellular loops (ECL1-3), three intracellular loops (ICL1-3), the N-terminus (N-Term) and C-terminus (C-Term). Ligand binding (blue) involves extracellular and transmembrane domains and consequently triggers conformational changes, which are transmitted to intracellular domains for induction of signaling cascades (green). **d:** Zoomed-in view on the multiple protein sequence alignment. Bovine rhodopsin (RHO) served as reference ^89^ to identify ligand binding and signaling domains of CHRM3-based DREADD, the human β2-adrenergic receptor (β2AR), three out of 292 GPCRs-of-interest (see **Supplementary Table S1**), human α1-adrenergic receptor (hα1AR) and hamster β2-adrenergic receptor (hamβ2AR). In grey: TMHMM-predicted transmembrane helices. In green: signaling domain for the first intracellular loop (ICL1). **e:** Crystal structures representing bovine RHO, rat CHRM3 as surrogate for DREADD, and human β2AR. Ligand binding domains (blue), signaling domains (green), and TMHMM-predicted sequences (grey) are highlighted. Structural representations are rotated with the intracellular domains facing the screen. Dotted lines: sequences not available within the crystal structures.

### Engineering chimeric DREADD-β2AR

Next, we designed our first CNO-inducible GPCR chimera DREADD-β2AR *in-silico* by combining DREADD ligand binding and β2AR signaling domains (**Fig.2a**). Additionally, we introduced two modifications to the N-terminus: a hemagglutinin-derived signal peptide ^51,52^ followed by a VSV-G epitope ^53^ to probe for cell surface expression. The signal peptide supports co-translational import into the endoplasmic reticulum and subsequent plasma membrane incorporation ^51,52^. Neither DREADD nor β2AR contain such a peptide sequence according to the SignalP algorithm ^54^ (**Supplementary Fig.S2a**). When we re-analyzed both GPCRs after introducing our N-terminal modifications *in-silico*, SignalP identified the signal peptide and its cleavage site upstream of the VSV-G tag (**Supplementary Fig.S2b**). We synthesized the DREADD-β2AR coding sequence and cloned it into a mammalian expression vector utilizing the ubiquitous CMV promoter. Then, we transfected HEK cells and after 24 hours performed immunostaining for the VSV-G tag under non-permeabilizing conditions. We confirmed that DREADD-β2AR successfully incorporated into the cell membrane based on the strong VSV-G signal (Fig.2b), whereas non-transfected HEK cells lacked this staining (**Supplementary Fig.S2c**). We also generated DREADD alone and non-chimeric β2AR containing the same N-terminal modifications; both showed successful surface expression (**Supplementary Fig.S2d-e**).

**Figure 2:**
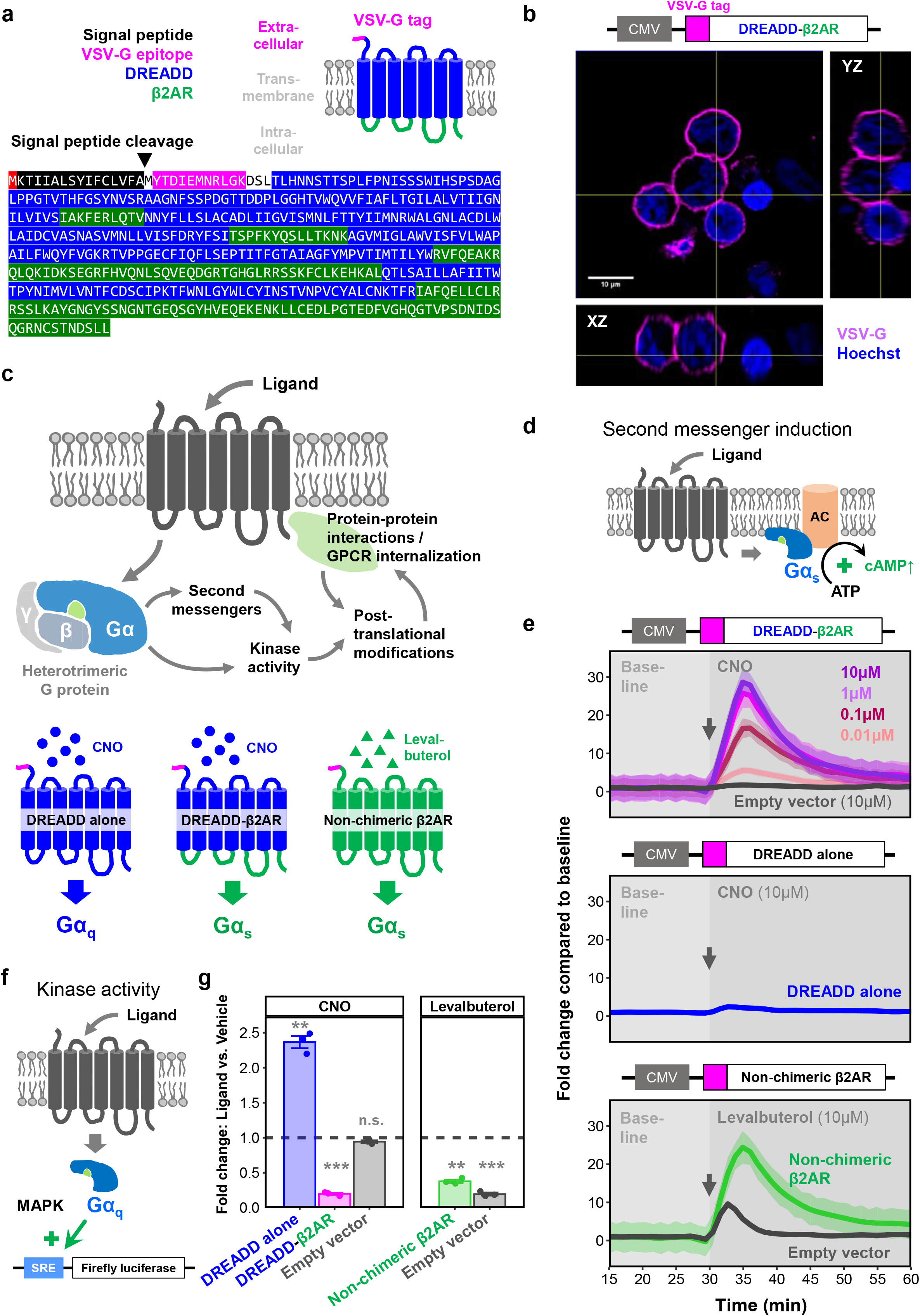
DREADD-β2AR recapitulates second messenger induction and MAPK activity of non-chimeric β2AR. **a:** Schematic of DREADD-β2AR and corresponding protein sequence encoding for signal peptide (black), VSV-G epitope (magenta), DREADD ligand binding domains (blue), and β2AR signaling domains (green). Black arrow: start of the mature GPCR after post-translational cleavage of the signal peptide. **b:** Orthogonal view of DREADD-β2AR-transfected HEK cells immunostained for the N-terminal VSV-G tag under non-permeabilizing conditions. Magenta: VSV-G tag. Blue: nuclear staining. CMV, human cytomegalovirus promoter. **c:** Schematic of signaling pathways for functional validation of the DREADD-based chimeras. The heterotrimeric G protein consists of an α- and βγ-subunit. Below: DREADD alone is a Gα_q_-coupled receptor, whereas non-chimeric β2AR recruits G proteins with a Gα_s_ subunit. DREADD-β2AR contains the β2AR signaling domains to recruit Gα_s_. **d:** Schematic of Gα_s_-coupled GPCR inducing cAMP synthesis after ligand stimulation through adenylyl cyclase (AC) activation. **e:** Real-time measurement of cAMP-dependent luciferase activity in HEK cells transfected with DREADD-β2AR (top), DREADD alone (middle), non-chimeric β2AR (bottom), or empty vector (top, bottom). Baseline measurements followed by ligand application (grey arrow for onset) of either CNO (top, middle) or levalbuterol (bottom). Graphs show fold changes compared to baseline mean. Ribbons: 95% confidence intervals of four repetitions. **f:** Schematic of Gα_q_-coupled GPCR engaging in the mitogen-activated protein kinase (MAPK) pathway which induces transcription of a firefly luciferase reporter from a serum responsive element (SRE). **g:** Endpoint measurement of SRE-dependent luciferase activity in HEK cells transfected with DREADD alone (blue), DREADD-β2AR (magenta), non-chimeric β2AR (green), or empty vector (grey). Ligand stimulation either with CNO (left) or levalbuterol (right). Dashed line: level of vehicle control. Error bars: standard error of the mean of three repetitions. One-sample T-test: *p\*\*\** < 0.001; *p\*\** < 0.01; *p^ns^* > 0.05.

### Functional validation of chimeric DREADD-β2AR

To validate DREADD-β2AR functionality, we investigated whether CNO stimulation mimics the signaling pathways of non-chimeric β2AR as outlined in **Figure 2c**. First, we focused on the induction of second messenger cascades. β2AR is classically known to recruit Gα_s_ upon ligand binding, resulting in an increase of cytoplasmic cAMP due to adenylyl cyclase (AC) activation ^55^ (**Figure 2d**). We co-transfected HEK cells with DREADD-β2AR and a modified firefly luciferase that increases luminescence in the presence of cAMP ^56^. Indeed, we found a CNO-dose-dependent increase in cAMP with DREADD-β2AR (**Figure 2e**), which was not observed in cells transfected with empty vector backbone, or with DREADD alone. For comparison, we transfected cells with non-chimeric β2AR and applied the selective β2AR-agonist levalbuterol ^57,58^. DREADD-β2AR and non-chimeric β2AR elicited similar fold-changes when stimulated with their respective ligand at a 10µM concentration, which we subsequently used for all further assays. As a note, HEK cells endogenously express β2AR ^59^, which explains the partial response of empty vector-transfected cells to levalbuterol. Our data suggests that DREADD-β2AR successfully recapitulated the Gα_s_-induced cAMP upregulation of non-chimeric β2AR. Importantly, CNO stimulation of the Gα_q_-coupled DREADD alone did not impact cAMP levels, indicating that the DREADD-β2AR response was mediated through properly identified β2AR signaling domains.

Next, we investigated kinase activity (Fig.2c). Gα_q_-coupled GPCRs trigger the mitogen-activated protein kinase (MAPK) pathway and induce transcription through a serum responsive element (SRE) ^60^. Therefore, we measured luciferase activity driven by an SRE reporter (Fig.2f). As expected, HEK cells transfected with DREADD alone increased luciferase activity 2.5-fold upon stimulation with CNO compared to vehicle (Fig.2g). We hypothesized that this effect would be absent in DREADD-β2AR-transfected HEK cells. Indeed, DREADD-β2AR did not increase luciferase activity; instead, the activity decreased more than 2-fold, which was similar to the non-chimeric β2AR response upon levalbuterol treatment. In empty vector-transfected cells, CNO had no impact on SRE-dependent reporter transcription, while levalbuterol reduced luciferase activity due to endogenous β2AR expression in HEK cells ^59^. The opposing responses with DREAD-β2AR and DREADD alone further substantiate the correct identification of β2AR signaling domains.

Next, we focused on protein-protein interactions and post-translational modifications regulated by β2AR signaling (Fig.2c). Ligand-activated β2AR also recruits β-arrestin 2, which creates a scaffold for attracting signaling kinases ^25^ and further plays a role in receptor internalization ^19^. To investigate the interaction between β-arrestin 2 and non-chimeric β2AR or DREADD-β2AR, we attached the complementary luciferase subunits LgBiT and SmBiT to their C-termini, respectively ^61–63^. Upon β-arrestin 2 recruitment, both subunits are brought into close proximity, resulting in a bioluminescent signal (Fig.3a). Levalbuterol stimulation of non-chimeric β2AR, as well as CNO stimulation of DREADD-β2AR, immediately increased bioluminescence compared to vehicle treatment (Fig. 3b). This indicates that DREADD-β2AR recapitulates the fast β-arrestin 2 recruitment observed with non-chimeric β2AR.

**Figure 3:**
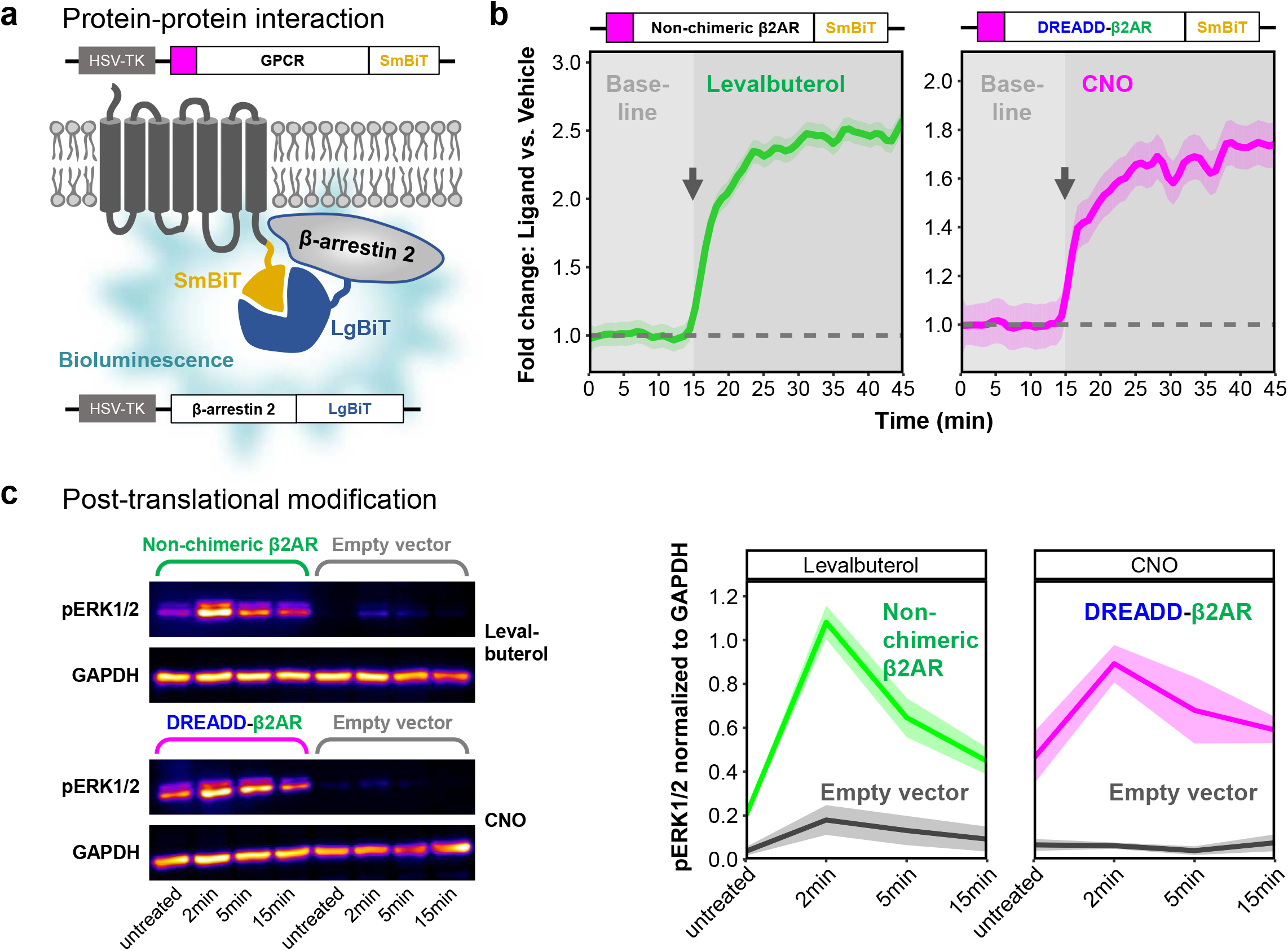
DREADD-β2AR recruits β-arrestin 2-and phosphorylates ERK1/2. **a:** Schematic of induced bioluminescence upon GPCR-SmBiT and β-arrestin 2-LgBiT interaction. GPCR, G protein-coupled receptor. HSV-TK, herpes simplex virus thymidine kinase promoter. **b:** Real-time measurement of bioluminescence in HEK cells transfected with non-chimeric β2AR (left) or DREADD-β2AR (right). Baseline measurements followed by ligand application (grey arrow shows onset) of either levalbuterol (left) or CNO (right). Graphs show baseline-normalized fold changes compared to vehicle. Dashed line: level of vehicle control. Ribbons: 95% confidence intervals of four repetitions. **c:** Phosphorylation analysis of extracellular signal-regulated kinases 1 and 2 (ERK1/2) in untreated, levalbuterol- or CNO-treated HEK cells transfected with non-chimeric β2AR, DREADD-β2AR, or empty vector. Left: Western blot for pERK1/2 and GAPDH (loading control). The anti-pERK1/2 antibody results in an upper band for pERK1 (44kDa) and a lower band for pERK2 (42kDA). Right: Densitometry analysis of combined pERK1/2 normalized to GAPDH. Ribbons: standard error of the mean of three repetitions.

β2AR signaling also involves the rapid phosphorylation of extracellular signal-regulated kinases 1 and 2 (ERK1/2), which is partly mediated through recruitment of β-arrestins ^25,55^. So, we investigated whether ERK1/2 phosphorylation occurred in HEK cells transfected with non-chimeric β2AR or DREADD-β2AR following treatment with levalbuterol or CNO, respectively. For both constructs, phosphorylation peaked two minutes after ligand stimulation and gradually declined after five minutes (Fig.3c, **Supplementary Fig.S3**), suggesting the recapitulation of post-translational modification dynamics. CNO exposure of empty vector-transfected HEK cells did not impact ERK1/2 phosphorylation.

β-arrestin recruitment also mediates GPCR internalization ^18–20^, which provides a regulatory feedback loop for receptor activity after ligand stimulation (Fig.2c) ^64,65^. To visualize receptor trafficking, we engineered DREADD-β2AR with EGFP attached at the C-terminus (Fig.4a). We transfected this construct into HEK cells and 24 hours later incubated them for 30 minutes with anti-VSV-G antibody to distinguish cell surface-incorporated DREADD-β2AR from receptors retained within the cell. Colocalization of VSV-G antibody and EGFP occurred on the cell surface. We barely found VSV-G signal within transfected cells suggesting that DREADD-β2AR internalization is absent without ligand stimulation (Fig.4b). In contrast, when we applied CNO for either 15, 30 or 60 minutes following VSV-G antibody labeling, VSV-G/EGFP signals colocalized within the cytoplasm. Internalization increased after 30 minutes and became significantly higher after 60 minutes of CNO exposure compared to vehicle treatment (Fig.4c-e). We conclude that DREADD-β2AR can undergo ligand-induced receptor internalization.

**Figure 4:**
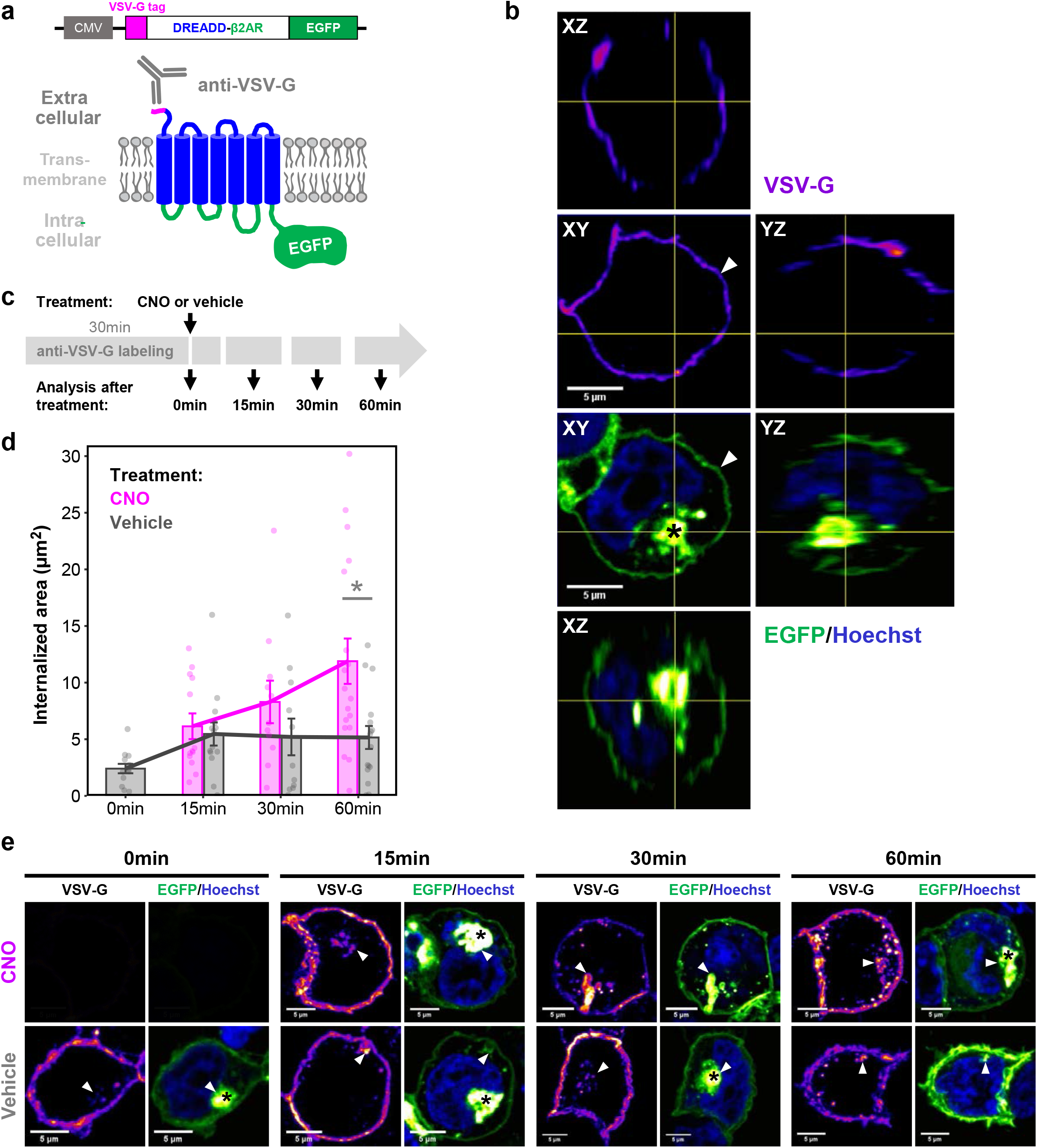
Internalization of DREADD-β2AR following CNO stimulation. **a:** Schematic of the DREADD-β2AR-EGFP construct for internalization analysis. C-terminal EGFP visualizes receptor trafficking within the cell. Cell surface-expressed receptors are labeled with an antibody against the VSV-G epitope. CMV, human cytomegalovirus promoter. **b:** Orthogonal view of DREADD-β2AR-EGFP-transfected HEK cell fixed immediately after 30 minutes of VSV-G antibody labeling. VSV-G signal visualized with an intensity-based color code (purple-red-yellow) to display signals of varying intensities. Green: EGFP visualized with intensity-based color code (green-white). Blue: nuclear staining with Hoechst. White arrow head: VSV-G/EGFP signal at the cell surface. Black asterisk: accumulation of cytoplasmic EGFP indicating receptors retained within the cell. **c:** Schematic of experimental design. Following 30 minutes of antibody incubation, cells were fixed either immediately (0 minutes) or 15, 30 and 60 minutes after exposure to CNO or vehicle. **d:** Stimulation of DREADD-β2AR-EGFP-transfected HEK cells with CNO (magenta) or vehicle (grey). Each dot shows internalized VSV-G-positive area of a single cell. Error bars: standard error of the mean within each condition. Magenta and grey lines connect the mean values of CNO or vehicle exposure times, respectively. Linear regression analysis: *p*\* < 0.05. **e:** Representative maximum intensity projections of individual cells analyzed for internalized receptors confirmed by colocalizing VSV-G/EGFP signal (white arrow heads).

Together, our results confirm that DREADD-β2AR successfully recapitulates the signaling cascades (Fig.2c) of non-chimeric β2AR with similar dynamics.

### Chimeric DREADD-β2AR recapitulates β2AR-mediated effects on microglia motility

Microglia are highly motile cells that constantly scan their environment for signs of disrupted tissue homeostasis. Activation of β2AR signaling was recently shown to rapidly induce filopodia formation as a consequence of elevated cAMP levels ^45^. Indeed, when we performed live-imaging of primary microglia cultures, we confirmed filopodia extension and an increase in total microglia area after levalbuterol application (**Fig.5**). During the first 10 minutes of baseline recordings, microglia were motile and changed their area only marginally. After levalbuterol stimulation, the cell area significantly increased throughout the following 45 minutes of imaging compared to the baseline (Fig.5a, e). To recapitulate this phenotype with our DREADD-β2AR, we first generated a bicistronic GPCR-P2A-EGFP vector containing a self-cleaving P2A peptide site ^66^ that allows simultaneous GPCR and cytoplasmic EGFP expression (**Supplementary Fig.S4a**). We transfected HEK cells with this DREADD-β2AR-P2A-EGFP vector and confirmed the expected cytoplasmic EGFP localization co-existing with anti-VSV-G immunostaining on the cell membrane (**Supplementary Fig.S4b**). Subsequently, we packaged our DREADD-β2AR-P2A-EGFP construct into lentiviral vectors and transduced primary microglia. When we imaged EGFP-positive cells, we found that CNO application induced filopodia formation (Fig.5b, e) similar to levalbuterol. Untransduced microglia stimulated with either vehicle or CNO did not significantly increase their area (Fig.5c-e), suggesting that DREADD-β2AR successfully mimics β2AR signaling in microglia and can modulate primary microglia function.

**Figure 5:**
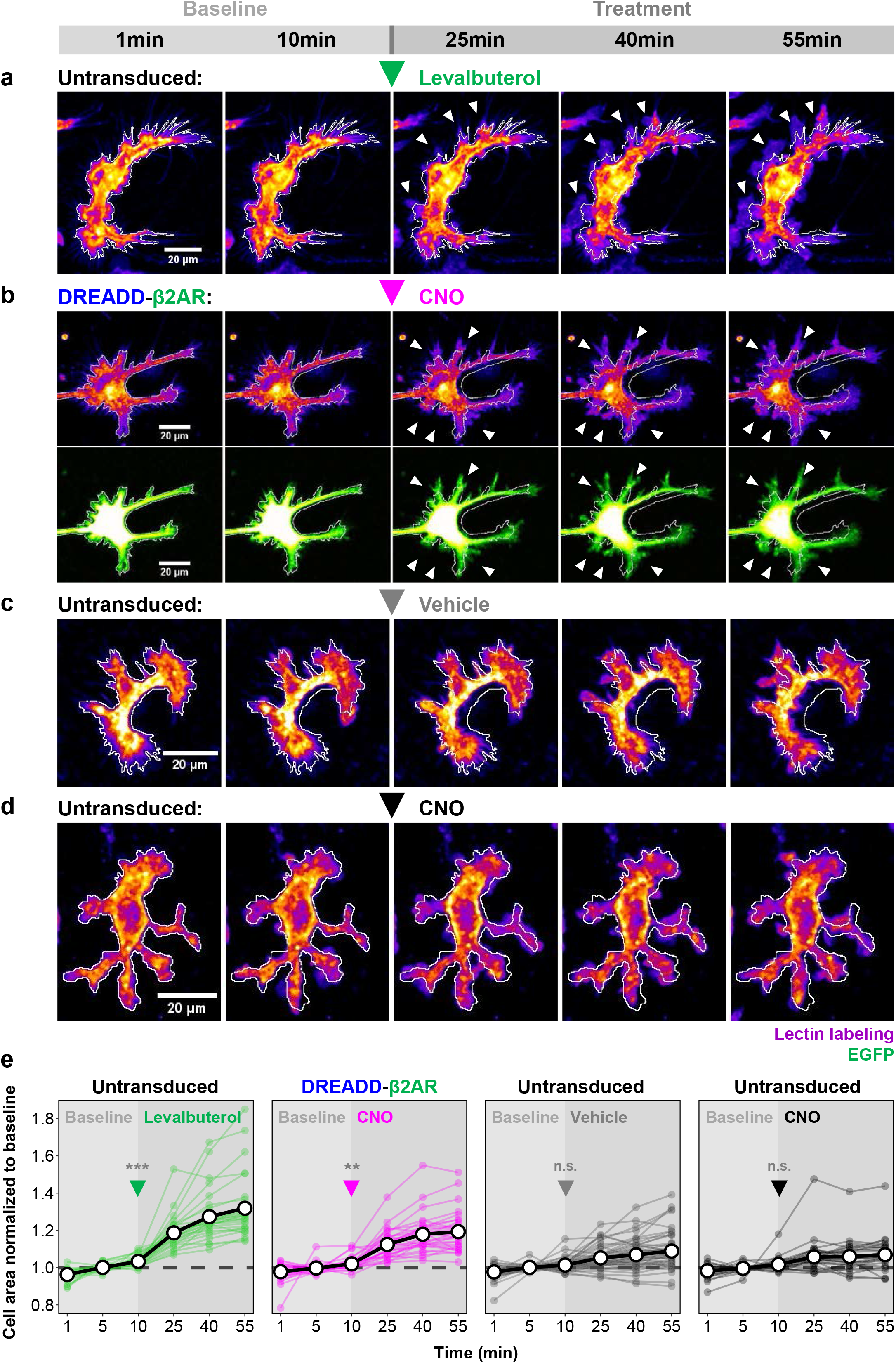
Stimulation of DREADD-β2AR induces filopodia formation in primary microglia. a-d: Representative images of primary microglia during 55 minutes of live imaging. After a 10 minutes baseline, untransduced cells were stimulated with either levalbuterol (**a**), vehicle (**c**), or CNO (**d**). Cells transduced with DREADD-β2AR-P2A-EGFP (**b**) were treated with CNO after 10 minutes of baseline recording. Lectin labeling visualized with an intensity-based color code (purple-red-yellow) to display signals of varying intensities. Green: EGFP visualized with intensity-based color code (green-white). White outline: cell area perimeter at 1 minute projected on all other shown time points. White arrow heads: filopodia formation. **e:** Quantification of cell area changes throughout 55 minutes of live imaging. A 10 minutes baseline was followed by ligand application (arrow heads for onset) of either levalbuterol, CNO, or vehicle. Graphs show the fold change of individual cells normalized to their baseline mean at selected time points of 1, 5, 10, 25, 40, and 55 minutes. Thick black lines: mean of all cells (30-50 per experimental group). Thin colored lines: individual cells. Dashed lines: baseline mean. Linear regression analysis: *p\*\*\** < *0.001*; *p\*\** < *0.01*; *p^ns^* > *0.05*.

### Generating DREADD-based chimeras for additional microglial GPCRs-of-interest

After confirming the functionality of our strategy with DREADD-β2AR, we decided to extend our approach to GPR65 and GPR109A/HCAR2, which like β2AR, showed microglia-enriched gene expression (Fig.1a). GPR65 and GPR109A respond to protons ^67^ and ketone bodies ^68^, respectively, and were shown to modulate inflammatory responses such as cytokine expression in microglia *in-vitro* systems ^69,70^. Both of their ligands are prone to cause off-target effects as acidic environments trigger various unpredictable responses in immune cells ^71,72^, and the ketone β-hydroxybutyrate can impact histone modification in a GPCR-independent manner ^73,74^. This makes GPR65 and GPR109A interesting candidates for DREADD-based chimeras to dissect their inflammatory role with a well-defined ligand. Thus, we designed DREADD-GPR65 and DREADD-GPR109A with the same N-terminal modifications as DREADD-β2AR (Fig.2a). First, we transfected HEK cells with these chimeras and confirmed successful cell membrane incorporation through immunostaining for the VSV-G tag (Fig.6a-b). Then, we investigated whether both chimeras triggered their expected second messenger cascades and kinase activity. Like β2AR, GPR65 belongs to the Gα_s_-coupling family ^67^. Therefore, we applied our previously established validation strategy for second messenger induction (Fig2.d-e). We measured cAMP levels in HEK cells transfected with DREADD-GPR65 and found a significant increase after CNO stimulation, which was not detected in empty vector-transfected cells (Fig.6c). DREADD-GPR65 also impacted the MAPK pathway and reduced SRE-mediated reporter expression, similar to β2AR (Fig.6d, Fig.2f-g).

**Figure 6:**
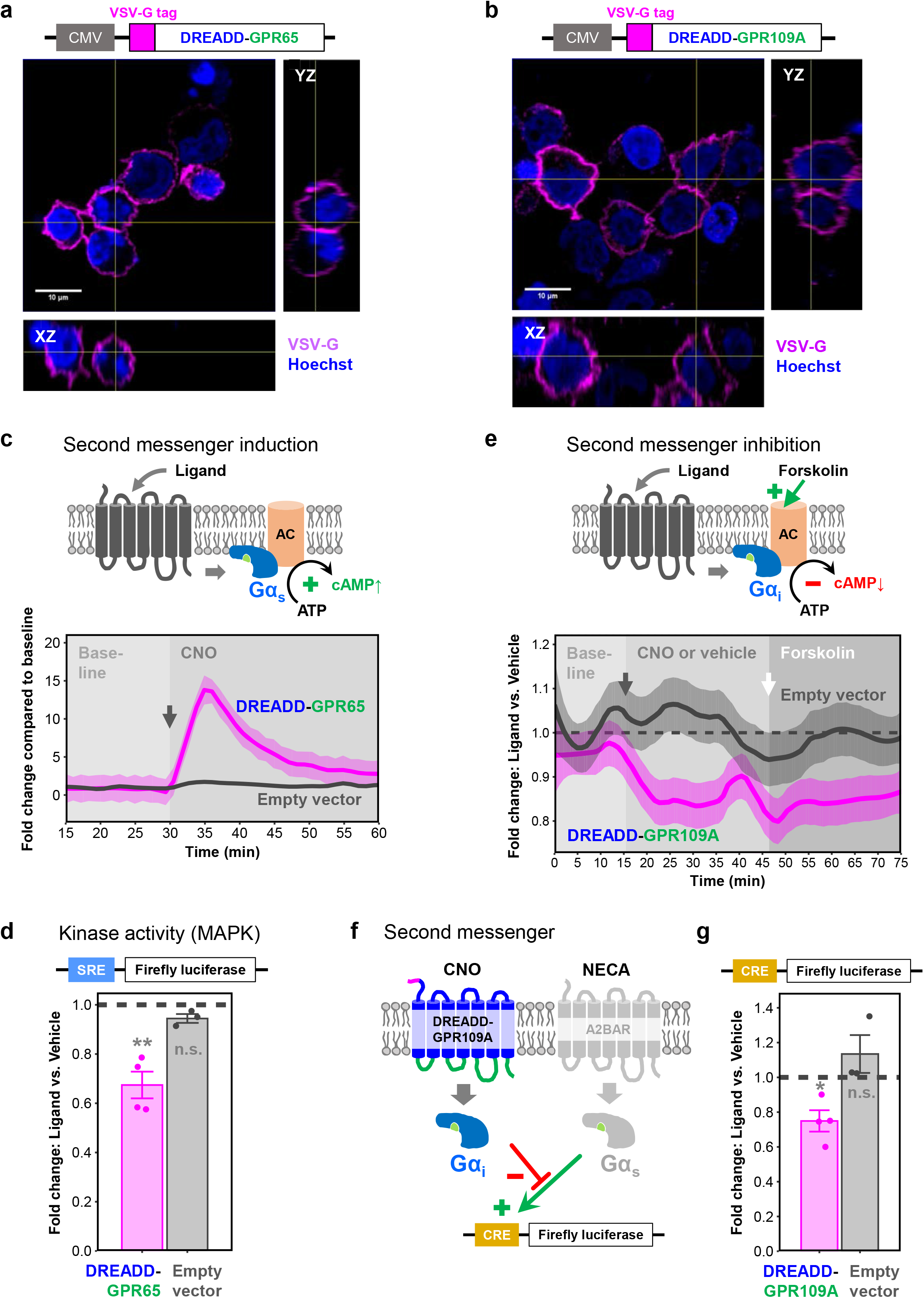
DREADD-GPR65 and DREADD-GPR109A respond with their expected signaling cascades. a-b: Orthogonal view of HEK cells transfected with DREADD-GPR65 (**a**) or DREADD-GPR109A (**b**) immunostained for the VSV-G tag under non-permeabilizing conditions. Magenta: VSV-G tag. Blue: nuclear staining with Hoechst. CMV, human cytomegalovirus promoter. **c:** Top: Schematic of Gα_s_-coupled GPCR inducing cAMP synthesis after ligand stimulation through adenylyl cyclase (AC) activation. Below: Real-time measurement of cAMP-dependent luciferase activity in HEK cells transfected with DREADD-GPR65 (magenta) or empty vector (grey). Baseline measurements followed by CNO application (grey arrow for onset). Graphs show fold changes compared to baseline mean. Ribbons: 95% confidence intervals of four repetitions. **d:** Endpoint measurement of serum responsive element (SRE)-dependent luciferase activity in HEK cells transfected with DREADD-GPR65 (magenta) or empty vector (grey). Ligand stimulation with CNO. Dashed line: level of the respective vehicle control. Error bars: standard error of the mean of three to four repetitions. One-sample T-test: *p\** < 0.05; *p^ns^* > 0.05. **e:** Top: Schematic of Gα_i_-coupled GPCR reducing cAMP levels after ligand stimulation through adenylyl cyclase (AC) inhibition. Forskolin induces cAMP synthesis through AC activation. Below: Real-time measurement of cAMP-dependent luciferase activity in HEK cells transfected with DREADD-GPR109A (magenta) or empty vector (grey). Baseline measurements followed by application of CNO or vehicle (grey arrow for onset) and forskolin (white arrow for onset). Graph shows fold changes compared to vehicle. Dashed line: level of the vehicle control. Ribbons: 95% confidence intervals of five repetitions. **f:** Schematic of competition assay between Gα_i_-coupled DREADD-GPR109A and Gα_s_-coupled A2B adenosine receptor (A2BAR). Simultaneous stimulation of Gα_i_ through CNO and Gα_s_ through NECA prevents cAMP responsive element (CRE)-mediated luciferase reporter activity. **g:** Endpoint measurement of CRE-dependent luciferase activity in HEK cells transfected with DREADD-GPR109A (magenta) or empty vector (grey). Ligand stimulation with CNO simultaneously with NECA. Dashed line: level of the respective vehicle control. Error bars: standard error of the mean of three to four repetitions. One-sample T-test: *p\** < 0.05; *p^ns^* > 0.05.

In contrast to GPR65 and β2AR, GPR109A couples to Gα_i_ and suppresses cAMP synthesis by inhibiting adenylyl cyclase (AC) ^68^ and therefore competes with the AC activator forskolin ^75,76^ (Fig.6e). To measure Gα_i_-mediated decreases in cAMP, we adapted a cAMP-dependent luciferase assay with kinetics suitable for Gα_i_-signaling ^77^. Within 10 minutes following CNO stimulation, DREADD-GPR109A-transfected HEK cells decreased cAMP levels by approximately 15% compared to vehicle (Fig.6e). After 30 minutes of CNO exposure, we added forskolin as a competing component to induce cAMP synthesis. DREADD-GPR109A-transfected cells exposed to CNO kept their cAMP signal approximately 15% below the vehicle control suggesting robust AC inhibition. Empty vector-transfected HEK cells did not respond to CNO, and their cAMP levels always remained at vehicle control levels (Fig.6e, **Supplementary Fig.S5a**).

To further substantiate the Gα_i_ effect, we tested for the ability of DREADD-GPR109A to compete with Gα_s_ signaling (Fig.6f). For this, we used a reporter that drives luciferase expression through a cAMP-responsive element (CRE) ^60^, which is induced by Gα_s_ activity. First, we confirmed successful Gα_s_ induction in empty vector-transfected HEK cells through stimulation with 5’-N-ethylcarboxamidoadenosine (NECA), a potent agonist of the endogenously expressed Gα_s_-coupled A2B adenosine receptor (A2BAR) ^78^. The expression of the CRE reporter was NECA dose-dependent and reached saturation at 5µM, while concomitant CNO application did not interfere (**Supplementary Fig.S5b**). Subsequently, we transfected HEK cells with DREADD-GPR109A and applied CNO together with 5µM NECA. As anticipated, CNO significantly inhibited Gα_s_-mediated transcription from the CRE reporter by approximately 20% when compared to vehicle (Fig.6g). In addition, DREADD-GPR109A had no impact on the MAPK pathway measured through SRE reporter activity (**Supplementary Fig.S5c**). Even though DREADD-GPR65 and DREADD-GPR109A differed in their second messenger and kinase activity, both GPCRs were able to recruit β-arrestin 2 emphasizing that individual GPCRs can display diverse signaling patterns (**Fig.S5d**). These results suggest that our DREADD-based strategy is reproducible and can be extended to other GPCRs-of-interest.

### DREADD-based chimeras modulate microglial gene expression under inflammatory conditions

Finally, we utilized our DREADD-based chimeras to investigate immunomodulatory consequences of GPCR signaling. We took advantage of the microglia-like cell line HMC3 ^79^, which has been used to study inflammation ^80^ and allows generation of cell lines with stable DREADD-GPCR expression. The latter is advantageous for reliable quantification of inflammatory responses and cannot be achieved in primary microglia due to suboptimal transduction efficiencies with available vectors ^81^. Thus, we cloned and packaged each DREADD-GPCR-P2A-EGFP construct into genome-integrating lentiviral vectors, transduced HMC3 cells, and fluorescence-activated cell sorted for EGFP-positive cells. We confirmed successful incorporation of GPCR chimeras in EGFP-expressing HMC3 cells through VSV-G immunostaining under non-permeabilizing conditions (**Supplementary Fig.S6a-c**).

Next, we induced inflammation by exposing HMC3 cells to recombinant interferon γ (IFNγ) and interleukin 1β (IL1β). Both cytokines can trigger the transcription of inflammatory genes like interleukin 6 (IL6) ^80^. Tumor necrosis factor (TNF) and IL1β expression are also part of the HMC3 cell inflammatory signature ^82^. Quantitative reverse transcription PCR (RT-qPCR) confirmed that IFNγ/IL1β stimulation increased the expression of these three inflammatory genes (**Supplementary Fig.S7a**). In parallel, we also confirmed that β2AR endogenously occurs in HMC3 cells at moderate mRNA levels (**Supplementary Fig.S7b**), allowing stimulation with the selective β2AR agonist levalbuterol. Thus, we treated non-transduced HMC3 cells with levalbuterol and compared the abundance of IL6, TNF, and IL1β transcripts to the mean of untreated samples. Neither of the three genes responded to levalbuterol treatment, whereas IFNγ/IL1β stimulation distinctly increased their expression. However, when we combined levalbuterol with IFNγ/IL1β, we found significant changes compared to IFNγ/IL1β stimulation alone. Transcript levels of IL6 increased and TNF decreased, while at the same time IL1β stayed unaltered (Fig.7a). Strikingly, we recapitulated the same response pattern with DREADD-β2AR-expressing cells upon CNO application (Fig.7b). Importantly, CNO did not impact the inflammatory response in the absence of GPCR chimeras. (Fig.7c). We repeated this experiment with the DREADD-GPR65 cell line and found a similar effect with increased IL6 and dampened TNF expression (Fig.7d). In contrast, DREADD-GPR109A did not significantly modify inflammatory gene expression induced by IFNγ/IL1β stimulation (Fig.7e).

**Figure 7:**
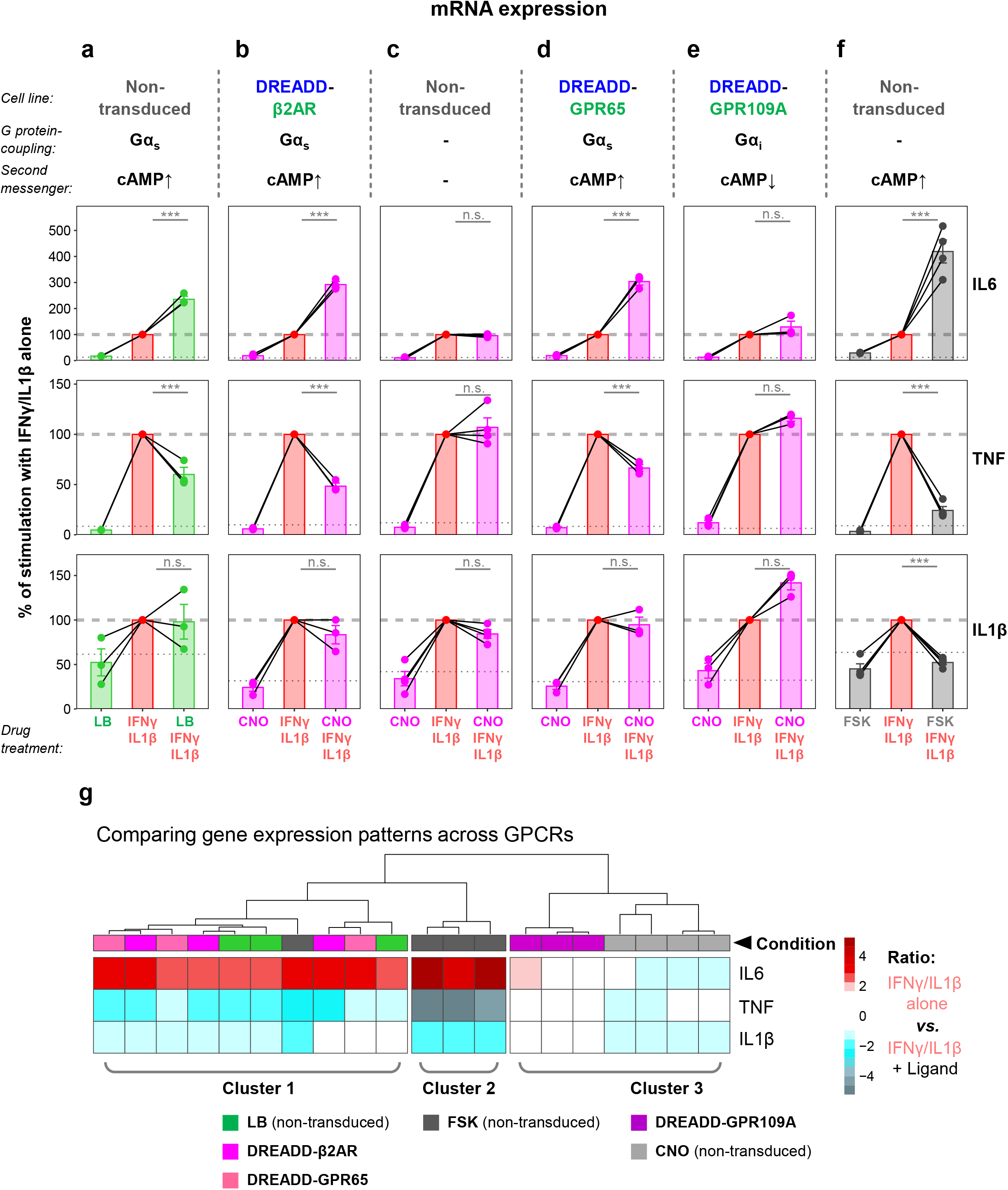
DREADD-based chimeras modulate inflammatory gene expression. a-f: Quantitative reverse transcription PCR (RT-qPCR) for interleukin 6 (IL6, top row), tumor necrosis factor (TNF, middle row), and interleukin 1β (IL1β, bottom row). Different HMC3 cell lines simultaneously treated with recombinant interferon γ (IFNγ) and interleukin 1β, and combinations of levalbuterol (LB, **a**), CNO (**b-e**), or forskolin (FSK, **f**). Graphs show fold changes compared to untreated cells with the IFNγ/IL1β treatment set to 100% within each repetition (dashed line). Dotted line: level of untreated controls. Lines connecting dots: dependent samples within experimental repetitions. Error bars: standard error of the mean of three to four repetitions. Linear regression analysis: *p\*\*\** < *0.001*; *p^ns^* > *0.05.* **g:** Hierarchical clustering of the gene expression pattern for IL6, TNF, and IL1β upon IFNγ/IL1β simulation. Conditions are color-coded corresponding to combinations of cell line and drug treatment as outlined in panels **a-f**.

Since all three Gα_s_-coupled receptors modulated gene expression in the same manner, and we have previously shown their ability to induce the second messenger cAMP (Fig.2e, Fig.6c), we hypothesized that elevated cAMP levels during IFNγ/IL1β exposure are responsible for the shared gene expression signature. To test this, we performed IFNγ/IL1β stimulation in the presence of forskolin, which induces cAMP synthesis in a GPCR-independent manner ^75,76^.

Indeed, we observed a significant increase of IL6 and decrease of TNF compared to IFNγ/IL1β treatment alone (Fig.7f). Interestingly, forskolin prevented the IFNγ/IL1β-mediated upregulation of IL1β mRNA levels, which we did not observe with endogenous β2AR, DREADD-β2AR, or DREADD-GPR65. When we applied hierarchical clustering to characterize the inflammatory gene expression pattern across different GPCRs, we confirmed that endogenous β2AR, DREADD-β2AR and DREADD-GPR65 intermingled within cluster 1 and were distinct from the forskolin treatment, which forms cluster 2 (Fig.7g). The third cluster contained DREADD-GPR109A and CNO-treated non-transduced cells, which both did not significantly modify the IFNγ/IL1β-induced response. These results suggest that our DREADD-based strategy provides the means to mimic GPCR signaling with high fidelity, which is not achieved solely by triggering the underlying second messenger cascade with forskolin.

## Discussion

Here, we illustrate the utility of a DREADD-based GPCR chimera strategy to selectively dissect the impact of GPCR activation in microglia. DREADD-based chimeras exploit the advantages of the DREADD system, which responds to CNO with high affinity and in a concentration range that minimizes potential off-target effects ^40–42^. This strategy complements existing light-inducible GPCR chimera approaches ^26–30^ and overcomes two main caveats associated with light stimulation such as phototoxicity ^32–35^ and the necessity for invasive surgical procedures for deep tissue light delivery ^36^. Not only can CNO be administered intraperitoneally and pass the blood brain barrier ^43^, but it also provides future opportunities to manipulate cells outside of light-accessible tissues such as circulating T-cells, B-cells, monocytes and granulocytes.

Even though our study focuses on immune cell function, GPCRs modulate a wide range of biological processes in other cell types as well. We generated a table with the protein sequences of putative signaling domains for all 292 GPCRs-of-interest included in our alignment, separated into ICL1-3 and C-terminus (**Supplementary Table S1**). These sequences can be inserted in-between the DREADD ligand binding domains as outlined in **Supplementary Figure S8** to generate a CNO-responsive chimera mimicking a GPCR-of-interest, which offers straightforward *in-silico* design of chimeric GPCRs.

### DREADD-based GPCR chimeras successfully mimic a GPCR-of-interest

Our engineering approach utilized multiple protein sequence alignment to identify CNO-binding DREADD domains and signaling domains of potential GPCRs-of-interest rather than protein domain identification on crystal structures; the latter are not available for most GPCRs. We used published crystal structures for RHO, CHRM3, and β2AR, and confirmed alignment accuracy by comparing our identified domains with these structural representations (Fig.1e). To validate our strategy, we focused on β2AR, given the extensive literature sources on its function and importance for the immune system ^9–11,44,45,49^. Indeed, we found that CNO stimulation of DREADD-β2AR in HEK cells successfully mimicked β2AR signaling and induced cAMP synthesis (Fig.2e), acted on the MAPK pathway by suppressing transcription from an SRE reporter (**Fig.2g**), recruited β-arrestin 2 (Fig.3b), and phosphorylated ERK1/2 (**Fig.3c**). This suggests that we properly identified ligand binding and signaling domains in DREADD and β2AR. Strikingly, we found that DREADD-β2AR-expressing primary microglia responded to CNO with filopodia formation, replicating previously reported effects of endogenous β2AR activation (**Fig.5**) ^45^. This underlines that our chimeric approach is able to modulate microglia function in primary culture systems.

### DREADD-based GPCR chimeras modulate microglia inflammatory gene expression

To dissect immunomodulatory properties of GPCR activation, we utilized the HMC3 microglia-like cell line and established cultures with stable DREADD-GPCR expression to reliably quantify the impact on inflammation. We challenged these cells with the inflammation mediators IFNγ and IL1β, which can induce prominent inflammatory gene expression in the HMC3 line ^80^ in contrast to commonly used lipopolysaccharide (LPS) ^83^. In the presence of IFNγ/IL1β, β2AR activation with levalbuterol induced pro- and anti-inflammatory properties reflected by enhanced IL6 and reduced TNF mRNA expression, respectively (**Fig.7a**). We successfully mimicked this response with DREADD-β2AR upon CNO stimulation (**Fig.7b**). These findings are in line with studies reporting similar pro- and anti-inflammatory effects of β2AR in different *in-vitro* systems ^44,84,85^. In our study, we also generated DREADD-based chimeras imitating the proton-sensing GPR65 and ketone-binding GPR109A. Following CNO stimulation, DREADD-GPR65 modified inflammatory gene expression similar to β2AR (**Fig.7d**), whereas DREADD-GPR109A did not alter IL6, TNF, or IL1β mRNA levels during IFNγ/IL1β-induced inflammation (**Fig.7e**). Our results in HMC3 cells during IFNγ/IL1β-induced inflammation highlights the response specificity with different *in vitro* systems and inflammation mediators ^69,70^.

Recently, the DREADD system has been explored for selective microglia manipulation *in-vivo* to study their role during neuropathic pain in mice ^86^. This study exploited a transgenic mouse line to achieve microglia-specific expression of a Ga_i_-coupled DREADD and to shed light on this broad signaling pathway. However, microglia might express Ga_i_-coupled receptors with non-canonical signaling cascades that are not captured by this DREADD approach. In this context, DREADD-based chimeras could offer a strategy for a more fine-tuned dissection of specific GPCRs and their role in regulating microglia function. While microglia *in-vitro* models are critical for neuroimmunological research, it is important to note that different culture systems display distinct genetic signatures and only partially reflect the phenotype of microglia *in-vivo* ^87^. Therefore, it would be ultimately desirable to apply DREADD-chimeras in an *in-vivo* context. However, *in-vivo* targeting of microglia is a major challenge within the field due to a current lack of efficient and specific vectors ^88^. Yet, GPCR signaling is critical for many other cell types that might be more accessible for chimeric GPCR expression. Our strategy complements existing methods for GPCR investigation and offers an alternative approach to dissect GPCR signaling in various contexts and model systems.

## Acknowledgements

We thank the scientific service units at IST Austria, in particular Flavia Gama Gomes Leite of the Molecular Biology Facility for excellent support with viral vector production, the Bioimaging Facility, and the Preclinical facility. We are grateful to the Novarino group for providing reagents and equipment, Harald Janoviak for sharing his expertise in synthetic biology, Marco Benevento for comments on the manuscript, and all members of the Siegert group for constant feedback on the project. This research was supported by a DOC Fellowship awarded to R.S. by the Austrian Academy of Sciences. We thank Life Science Editors for editing assistance.

## Author contributions

Conceptualization and Methodology: R.S. and S.S.; Formal Analysis: R.S.; Validation and Investigation: R.S., M.K-D, G.C.; Writing of Original Draft and Visualization: R.S. and S.S.; Supervision: S.S.

## Methods

### Animals

C57BL/6J (Cat#000664) were purchased from The Jackson Laboratories. All mice were housed in the IST Austria Preclinical Facility with a 12 hour light-dark cycle, food and water provided *ad libitum*. All animal procedures are approved by the Bundesministerium für Wissenschaft, Forschung und Wirtschaft (bmwfw) Tierversuchsgesetz 2012 (TVG 2012), BGBI. I Nr. 114/2012, idF BGBI. I Nr. 31/2018 under the numbers 66.018/0005-WF/V/3b/2016, 66.018/0010-WF/V/3b/2017, 66.018/0025-WF/V/3b/2017, 66.018/0001_V/3b/2019, 2020-0.272.234.

### Multiple protein sequence alignment

The previously established domains of bovine rhodopsin (RHO) ^89^ served as reference for the identification of ligand binding and signaling domains. In total, 294 protein sequences were aligned, including RHO, DREADD, two sequences (hamβ2AR, hα1AR) from Airan *et al.* ^89^ as internal control, and GPCRs-of-interest (human and mouse class A GPCRs available at IUPHAR/BPS; www.guidetopharmacology.org). Sequences were combined in a FASTA file, which served as input for the alignment algorithm MUSCLE ^90^. To visualize results, the alignment output was imported into the software Jalview 2.9.0b2. Sequences were identified as ligand binding or signaling domains based on their alignment with the RHO reference. Signaling domains were labeled according to their location as intracellular loops (ICL) 1-3 and C-terminus (C-Term).

### Predicting transmembrane GPCR domains

The bioinformatics tool TMHMM (www.cbs.dtu.dk/services/TMHMM) ^50^ was used to predict transmembrane helices (TMs) for selected GPCRs (RHO, DREADD, hamβ2AR, hα1AR, β2AR, GPR65, GPR109A, and GPR183). We highlighted predicted TMs in our alignment output with Jalview 2.9.0b2 (**Fig.1d**, **Supplementary Fig.S1**).

### Identifying GPCR domains on available crystal structures

We accessed the PDB data base (www.rcsb.org) to download structural representations of bovine RHO, rat CHRM3 as surrogate for the DREADD, and human β2AR (PDB IDs: 1U19, 4U15 and 2RH1, respectively). Structural data were imported into the software VMD 1.9.2 and oriented with the intracellular domains facing towards the screen. We then highlighted alignment-identified ICL1-3, C-Term, and TMHMM-predicted TMs to see whether they map to their expected locations. In case of partly missing structural data, we used dotted lines as representation.

### Adding N-terminal modifications to GPCR chimeras

The bioinformatics tool SignalP (www.cbs.dtu.dk/services/SignalP) ^54^ was used to predict whether DREADD and β2AR contains a signal peptide. Since such sequence was not found, we added a hemagglutinin-derived signal peptide (KTIIALSYIFCLVFA) at the N-terminus ^51,52^. Additionally, we also included a VSV-G epitope (YTDIEMNRLGK) followed by a DSL linker immediately after the signal peptide ^53^ (**Supplementary Fig.S2a-b**). In the corresponding DNA sequences, the start codon (ATG) was removed to prevent leaky scanning and to ensure that all proteins contain the VSV-G tag.

### Obtaining DNA sequences for GPCR chimeras via gene synthesis

To generate a chimera for a GPCR-of-interest, we combined ligand-binding DREADD domains and GPCR-of-interest signaling domains *in-silico* (**Supplementary Fig.S8**, **Supplementary Table S1**). We identified the corresponding DNA sequences of these domains through the NCBI Consensus Coding Sequence (CCDS) data base ^91^ and added our N-terminal modifications. The entire coding sequence was then synthesized (www.eurofinsgenomics.eu) in the pEX-K4 vector. During synthesis, recognition sites for restriction enzymes (EcoRI or NotI, BamHI) were added up- and down-stream of the chimera. The same strategy was used to obtain N-terminally modified non-chimeric β2AR and DREADD.

### Cloning

For HEK cell assays, if not otherwise stated, GPCRs were excised from pEX-K4 and inserted into the mammalian expression vector pcDNA3.1(-) using EcoRI or NotI, and BamHI. To study protein-protein interactions, we utilized the NanoBiT system (Promega; N2014). GPCRs were amplified from pEX-K4 with primers carrying restriction sites for NheI and EcoRI. These restriction sites were then used to clone GPCRs into pBiT2.1-C[TK-SmBiT] in order to obtain GPCR-SmBiT fusion constructs. β-arrestin 2 was amplifying from pCDNA3.1(+)-CMV-bArrestin2-TEV (Addgene #107245) with Gibson Assembly primers compatible with the NEBuilder HiFi DNA Assembly Kit (New England BioLabs; E2621). β-arrestin 2 was subsequently assembled into pBiT1.1-C[TK-LgBiT], linearized by NheI and XhoI, in order to obtain β-arrestin 2-LgBiT.

To generate DREADD-β2AR-EGFP, we amplified DREADD-β2AR from pEX-K4, and EGFP from PL-SIN-PGK-EGFP (Addgene #21316) with Gibson Assembly primers. Both fragments were then assembled into pcDNA3.1(-), linearized by NotI and BamHI. Bicistronic constructs encoding for GPCR-P2A-EGFP were obtained through a cloning step involving an intermediate vector, encoding for mCherry-P2A-EGFP, which was previously generated in the laboratory. First, GPCRs were amplified from pEX-K4 with Gibson

Assembly primers and assembled into the intermediate vector, linearized by NheI and BamHI in order to excise mCherry and replace it with GPCRs. Finally, GPCR-P2A-EGFP was amplified from these vector intermediates with Gibson Assembly primers and assembled into pcDNA3.1(-), linearized by NotI and BamHI.

For lentivirus production, we used a modified transfer vector based on PL-SIN-PGK-EGFP. This plasmid was modified through Gibson Assembly by introducing a WPRE sequence downstream of EGFP followed by a microRNA9 sponge (miR9T), which was previously described for optimized microglia transduction ^92,93^. WPRE was amplified from pAAV-hSyn-tdTomato (a gift from the Jonas group at IST Austria). The miR9T sequence was synthesized (www.eurofinsgenomics.eu) in the pEX-A258 vector and subsequently amplified. Both fragments were then assembled into PCR-linearized PL-SIN-PGK-EGFP, which generated PL-SIN-PGK-EGFP-WPRE-miR9T. Finally, GPCR-P2A-EGFP was amplified with Gibson Assembly primers from the previously established pcDNA3.1(-) vectors and assembled into PL-SIN-PGK-EGFP-WPRE-miR9T, linearized by PstI and BsrGI.

### Cell lines

HEK293T cells were obtained from ATCC (CRL-3216) and cultured in HEK-complete medium, containing DMEM (ThermoFisher; 31966; with high glucose content, GlutaMAX and pyruvate), 10% (v/v) fetal bovine serum (FBS; Sigma; 12103C; heat-inactivated for 30 minutes at 56°C), 1% (v/v) non-essential amino acids (Sigma; M7145) and 1% (v/v) penicillin-streptomycin (ThermoFisher; 15140-122). Medium was sterile filtered (0.22µm; TPP; 99505) and stored at 4°C.

HMC3 cells were obtained from ATCC (CRL-3304) and cultured in EMEM-complete medium, containing EMEM (ATCC, 30-2003), 10% (v/v) FBS and 1% (v/v) penicillin-streptomycin. Medium was sterile filtered (0.22µm) and stored at 4°C.

### Cell maintenance

HEK cells were maintained in T75 flasks with 15ml medium. Culture conditions were 37°C and 5% CO_2_. In order to passage cells, old medium was aspirated and cell layer was washed with 10ml DPBS (37°C). PBS was aspirated and 3ml Trypsin-EDTA (ThermoFisher; 25300-054; 37°C) were added for approximately 1 minute until the cell layer detached.

Trypsinization was stopped with 10ml medium (37°C). Cells were pelleted at Trypsin-EDTA (ThermoFisher; 25300-054; 37°C). Supernatant was aspirated and pelleted cells were resuspended thoroughly in 10ml medium (37°C). Cells were counted and 0.5-0.75 million cells were transferred to a new culture flask within a final volume of 15ml medium (37°C). Cells were passaged every 3-4 days when they reached approximately 80% confluency. HMC3 cells were maintained in 10cm dishes (Sigma, CLS430167) with 10ml medium. Culture conditions were 37°C and 5% CO_2_. For passaging, old medium was aspirated and cell layer was washed with 10ml DPBS (37°C). PBS was aspirated and 3ml Trypsin-EDTA were added for approximately 5-15 minutes until the cell layer detached. Trypsinization was stopped with 10ml medium (37°C). Cells were counted from this suspension and 0.25-0.5 million cells were transferred to a new culture dish within a final volume of 10ml medium (37°C). Cells were passaged every 3-4 days when they reached approximately 80% confluency.

### Coating plates for HEK cell assays and immunostaining

White clear-bottom 96-well plates (Greiner Bio-One; 655098) and 8-well chamber slides (ibidi; 80826; growth area: 1cm^2^) were coated with 50µl and 100µl poly-L-ornithine (ready-to-use 0.1% (w/v) solution; Sigma; P4957) respectively. After 1 hour incubation at room temperature, wells were washed three times with 100µl sterile Milli-Q water and left to dry with an open lid for 1 hour under UV irradiation in a sterile laminar flow hood. Culture dished were then wrapped with Parafilm and stored at 4°C.

### HEK cell transfection

Cells were transfected by seeding them into wells of indicated culture vessels containing transfection mix. To avoid toxicity of antibiotics during transfection, cell suspensions were prepared in HEK-complete medium without penicillin-streptomycin. Polyethylenimine (PEI; Polysciences; 24765) was used as transfection reagent. A stock solution (1mg/ml) was prepared by dissolving PEI in Milli-Q water and adjusting the pH to 7. Aliquots were stored at -20°C. To make the transfection mix, plasmids were first diluted in Optimem (ThermoFisher; 51985034) to a total concentration of 40ng/µl. In parallel, PEI stock was diluted 1:10 in Optimem and incubated for 5 minutes at room temperature. Plasmid and diluted PEI stock were then mixed 1:1 to generate the transfection mix (containing 2.5µl PEI stock per µg DNA). After 20 minutes incubation at room temperature, this transfection mix was pipetted into wells followed by adding the desired number of cells (Supplementary Table S2). Assays were performed 24 hours after transfection.

### Confocal microscopy

Images of immunostainings were acquired on inverted Zeiss LSM800 or Zeiss LSM880 microscopes with either a 63x oil immersion or 20x air objective. Live imaging of primary microglia was performed on an inverted Zeiss LSM800 using a 20x air objective.

### Preparation of Antifade mounting medium

Mowiol 4-88 (2.4g; Sigma; 81381) and glycerol (4.8ml; Sigma; G7757) were combined with 6ml Milli-Q water and 12ml Tris buffer (0.2M; pH 8) and stirred overnight at room temperature. After letting the solution rest for 2 hours, it was incubated for 10min at 50°C in a water bath and then centrifuged at 4700 x g for 15min. The supernatant was combined with DABCO (Sigma; D27802) at 2.5% (w/v). Aliquots were stored at -20°C.

### VSV-G immunostaining for confirming cell surface expression of GPCRs

HEK cells were transfected with GPCR (600ng) in coated 8-well chamber slides (ibidi; 80826) as described above. After 24 hours, live cells were immunostained under non-permeabilizing conditions. For this, mouse monoclonal anti-VSV-G antibody conjugated to Cy3 (Sigma; C7706; LOT: 049M4837V) was first subjected to ultrafiltration to reduce the concentration of cytotoxic NaN_3_. The required amount of antibody was diluted in PBS (5ml) and applied to a Vivaspin 6 concentrator (Sartorius; VS0601; MWCO: 10kDa; 5ml volume). After centrifugation (4000 x g; 10 minutes; 4°C), the concentrate was diluted in cold cell culture medium to obtain a final 1:250 dilution of antibody. To start the immunostaining, medium was aspirated from the chamber slides and replaced with cold antibody-containing medium. Cells were incubated on ice for 1 hour. Wells were then washed three times with cold medium without antibody for 3 minutes each, followed by fixation with cold 4% (w/v) PFA in PBS for 15 minutes. After fixing, cells were washed once with PBS for 3 minutes and subjected to nuclear staining with Hoechst (New England BioLabs; 4082S; diluted 1:5000 in PBS) for 5 minutes. Wells were briefly washed with PBS before adding 200µl Antifade mounting medium. Chamber slides were stored at 4°C until imaging. All incubation steps were carried out with 200µl of the respective solution and under protection from light to avoid bleaching of the Cy3 fluorophore.

Confocal images were acquired with a 63x oil immersion objective. Images were processed in Fiji 1.51f by applying a gamma correction of 0.5 for better visualization of faint VSV-G signals, followed by a rolling ball background subtraction and a 2x2x2 median filter. For VSV-G staining of HMC3 cells, uncoated 8-well chamber slides were used and 3,500 cells were seeded per well together with GPCR-encoding lentivirus at a multiplicity of infection (MOI) of 5. The previously described staining procedure was performed after three to four days when cells reached approximately 80% confluency. Confocal images were acquired with a 20x air objective. Images were processed with 0.5 gamma correction and 2x2x2 median filtering.

### Real-time measurement of cAMP levels

All real-time luciferase assays were performed in CO_2_-independent medium (Leibovitz’s L15 (ThermoFisher; 21083027; no phenol red), 10% (v/v) FBS and 1% (w/v) penicillin-streptomycin). The medium was sterile filtered (0.22µm) and stored at 4°C. To measure Gα_s_-induced increases in cAMP, a cAMP-dependent firefly luciferase (GloSensor) ^56^ was used (the plasmid was a gift from the Janovjak group formerly at IST Austria). As luciferase substrate, a 100mM stock solution of beetle luciferin (Promega; E1602) was prepared in 10mM HEPES and stored at -20°C protected from light. HEK cells were transfected in a 96-well plate with GPCR or empty vector backbone (100ng), and GloSensor (100ng). After 24 hours, medium was replaced with 90µl CO_2_-independent medium (37°C) containing 2.22mM beetle luciferin (1:45 dilution of stock). The plate was then incubated for 15 minutes at 37°C in an incubator with atmospheric CO_2_ and then transferred to a plate reader (BioTek; Synergy H1) with the lid removed. Total bioluminescence was measured in each well every 1-2.5 minutes (37°C, 1 second integration time; 200 gain) for 30 minutes to establish a baseline. After the last baseline measurement, the plate was ejected and a multichannel pipette was used to quickly apply 10µl levalbuterol or CNO (prepared in CO_2_-independent medium; 10 times more concentrated than the desired concentration of either 0.01µM, 0.1µM, 1µM, or 10µM). The measurement was immediately continued for 1 hour. Individual experiments were always carried out in triplicates for each condition. For each well, a fold change in bioluminescence was calculated by normalizing luminescence to the mean of the baseline measurement. For final analysis, baseline-normalized values were pooled from individual experimental repetitions.

To measure Gα_i_-induced decreases in cAMP, we used a different cAMP-dependent firefly luciferase suitable for Gα_i_ signaling (GloSensor-22F; Promega; E2301) ^77^. As luciferase substrate, a 100X stock solution of cAMP reagent (Promega; E1290) was prepared in 10mM HEPES and stored at -80°C. HEK cells were transfected with GPCR or empty vector (100ng), and GloSensor-22F (100ng). After 24 hours, medium was changed to 80µl CO_2_-independent medium (37°C) containing 2% (v/v) cAMP reagent (1:50 dilution of stock). The plate was incubated for 2 hours at room temperature and atmospheric CO_2_ before starting the measurement in the plate reader as previously described. After a 15 minutes baseline, each well received 10µl CNO (prepared in CO_2_-independent medium; 10 times more concentrated than the desired concentration of 10µM), or an equal amount of vehicle (medium only). The measurement was immediately continued for 30 minutes. Then, 10µl forskolin (prepared in CO_2_-independent medium; 100µM) was added to each well for a final concentration of 10µM. The measurement immediately continued for another 30 minutes. Vehicle controls always received the same transfection mix as their corresponding treated condition. Individual experiment were always carried out in triplicates for each condition. For each experiment, fold changes were calculated by normalizing ligand-treated conditions to the respective vehicle controls. To do this, luminescence values at each time point were divided by the mean of the respective vehicle control triplicate. For final analysis, vehicle control-normalized values were pooled from different experimental repetitions.

### Real-time measurement of β-arrestin 2 recruitment

As luciferase substrate, the Nano-Glo Live Cell Substrate (Promega; N2011) was used. HEK cells were transfected in a 96-well plate with GPCR-SmBiT (100ng) and β-arrestin 2-LgBiT (100ng). After 24 hours, medium was replaced with 90µl CO_2_-independent medium (37°C) containing 1% (v/v) Nano-Glo Live Cell Substrate (1:100 total dilution; added from a freshly prepared 1:20 pre-dilution in LCS Dilution Buffer supplied with the reagent). The plate was then incubated for 10 minutes at room temperature and subsequently transferred to a plate reader (BioTek; Synergy H1) with the lid removed. Total bioluminescence was measured in each well every 40 seconds (room temperature, 1 second integration time; 200 gain) for 15 minutes to establish a baseline. After the last baseline measurement, the plate was ejected and a multichannel pipette was used to quickly apply 10µl levalbuterol or CNO (prepared in CO_2_-independent medium; 10 times more concentrated than the desired concentration of 10µM), or an equal amount of vehicle (medium only). The measurement immediately continued for 45 minutes. Vehicle controls always received the same transfection mix as their corresponding treated condition. Individual experiments were always carried out in triplicate for each condition. For each well, a fold change in bioluminescence was calculated by normalizing luminescence to the mean of the baseline measurement. These values were further used to obtain fold changes by normalizing ligand-treated conditions to the respective vehicle controls. To do this, baseline-normalized values at each time point were divided by the mean of the respective vehicle control triplicate. For final analysis, these values were pooled from different experimental repetitions.

### SRE reporter assay

Transcription-based luciferase reporter assays were performed with the Dual-Glo kit (Promega; E2920). HEK cells were transfected in a 96-well plate with GPCR or empty vector (95ng), SRE-dependent firefly luciferase (95ng; Promega; E1340) ^60^, and ubiquitously expressed renilla luciferase (9.5ng) to normalize for inter-assay variability (renilla luciferase was inserted into pcDNA3.1(-) and provided as a gift from the Janovjak group formerly at IST Austria). After 24 hours, HEK-complete medium was replaced with 90µl fresh HEK-complete medium. Each well was then treated with 10µl levalbuterol or CNO (prepared in HEK-complete medium; 10 times more concentrated than the desired concentration of 10µM), or an equal amount of vehicle (medium only). After 6 hours incubation at 37°C and 5% CO_2_, 50µl of medium were removed from each well and replaced by 50µl Dual-Glo luciferase reagent. Following a 10 minute incubation at room temperature, the plate was transferred to a plate reader to measure firefly luminescence (BioTek; Synergy H1; room temperature, 1 second integration time; 200 gain). Then, 50µl Stop&Glo reagent was added to each well, followed by another 10 minute incubation at room temperature, renilla luminescence was measured with the same parameters. For each well, a ratio was obtained by normalizing firefly luminescence to renilla luminescence as a control for transfection efficiency, cell number and enzyme activity. Vehicle controls always received the same transfection mix as their corresponding treated condition. Individual experiments were always carried out in triplicates for each condition. For each experiment, fold changes between ligand-treated conditions and the respective vehicle controls were obtained by dividing firefly-renilla ratios by the mean of the respective vehicle control triplicate. These values were pooled from different experimental repetitions for final analysis.

### CRE reporter assay

HEK cells were transfected as previously described for the SRE reporter assay with the exception of substituting the SRE reporter with CRE-dependent firefly luciferase ^60^ (Promega; E8471).

To quantify inhibition of Gα_s_ activity, HEK-complete medium was replaced after 24 hours with 80µl fresh HEK-complete medium. Each well was then treated with 10µl CNO (prepared in HEK-complete medium; 10 times more concentrated than the desired concentration of 10µM), or an equal amount of vehicle (medium only). Immediately afterwards, 10µl 5’-N-ethylcarboxamidoadenosine (NECA; prepared in HEK-complete medium; 50µM) was added to all wells for a final concentration of 5µM. After 6 hours incubation at 37°C and 5% CO_2_, luminescence was detected and data were analyzed as previously described for the SRE reporter assay.

### Western blot to quantify ERK1/2 phosphorylation

HEK cells were transfected in 6-well plates with GPCR or empty vector (6µg). After 24 hours, cells were serum starved for 4 hours by replacing medium with 1.9ml HEK-complete medium without FBS. Then, cells were treated with 100µl levalbuterol or CNO (prepared in HEK-complete medium without FBS; 20 times more concentrated than the desired concentration of 10µM). Control conditions were left untreated. Treated cells were harvested for protein isolation 2, 5 or 15 minutes after addition of ligand (cells were kept at 37°C and 5% CO_2_ during ligand exposure). Untreated cells were harvested immediately. Cells were harvested by placing the plate on ice, aspirating medium and adding 200µl ice cold and freshly prepared lysis buffer (50mM Tris pH 7.4, 300mM NaCl, 1mM EDTA, 1mM Na3VO4, 1mM NaF, 10% (v/v) Glycerol, 1% (v/v) IGEPAL CA-630, 1% (v/v) Protease inhibitor mix set 1 (Calbiochem; 539131); 1 Phosstop tablet (Sigma; 4906845001) per 10ml) to each well. Cells were detached with a cell scraper, transferred to 1.5ml microcentrifuge tubes and sonicated (15 seconds; room temperature; inside a water bath). Samples were then centrifuged (14,000 x g, 20 minutes, 4°C) and supernatants were transferred to fresh tubes. A small volume of each sample was used to immediately measure protein concentration with the Pierce BCA Protein Assay kit (ThermoFisher; 23227). The rest was combined with 6X loading dye (375mM Tris pH 6.8, 9% (w/v) SDS, 30% (w/v) glycerol, 0.06% (w/v) Bromophenol blue, 600mM DTT), cooked for 5 minutes at 95°C and stored at -20°C for subsequent Western blot analysis. Three individual experiments were performed with one well per condition in each. SDS-PAGE was performed by loading 10µg protein (approximately 10µl) on 8% acrylamide gels with running buffer containing 25mM Tris, 192mM glycine, and 0.1% (w/v) SDS. Electrophoresis was started at 90V constant (two gels per chamber) until samples transitioned from stacking to running gel. Electrophoresis continued at 110V constant until the 25kDa band of the marker left the gel (approximately 2 hours). Proteins were transferred to PVDF membranes (Sigma; IPFL00005) via tank blotting (300mA constant; 2 hours; 4°C; additional cooling insert) with transfer buffer containing 25mM Tris, 192mM glycine, and 20% (v/v) methanol. Successful transfer was briefly checked with Ponceau staining (0.1% (w/v) Ponceau S, 5% (v/v) acetic acid). Membranes were cut to include proteins ranging from 32-80kDa and blocked with 5% (w/v) BSA in TBST (20mM Tris, 150mM NaCl, 0.1% (v/v) Tween 20) for 1 hour at room temperature. Membranes were then incubated overnight at 4°C with rabbit anti-phosphorylated ERK1/2 antibody (Cell Signaling Technology; 9101S; LOT: 30; 1:1,000) in TBST containing 5% (w/v) BSA. Next day, membranes were washed three times with TBST for 10 minutes each and exposed to donkey anti-rabbit secondary antibody conjugated to horse radish peroxidase (GE Healthcare; NA934V; LOT: 16976257; 1:10,000) in TBST containing 5% (w/v) BSA for 2 hours at room temperature. Membranes were again washed three times with TBST followed by signal detection with either SuperSignal West Pico PLUS (Thermo Fisher; 34579) or SuperSignal West Femto (Thermo Fisher; 34094) and imaging (Amersham 600; GE Healthcare). Membranes were then stripped (pH 2.2, 0.2M glycine) for 30 minutes at room temperature, washed three times and blocked again followed by incubation with rabbit anti-GAPDH antibody (Sigma; ABS16; LOT: 3275069; 1:1,000; overnight; 4°C).

Membranes were washed three times and subjected to secondary antibody using our standard procedure. The membranes were again washed and GAPDH signal was detected and imaged. Densitometry of bands (pERK1, pERK2, GADPH) was performed with Bio-Rad Image Lab 6.0.1. Densities of pERK1 and pERK2 were then summed to generate a single value (pERK1/2). To normalize for protein loading variability, each pERK1/2 value was divided by the respective GAPDH band on the same membranes after striping.

### GPCR internalization assay

HEK cells were transfected with DREADD-β2AR-EGFP (600ng) in coated 8-well chamber slides (ibidi; 80826) as described above. After 24 hours, live cells were subjected to anti-VSV-G antibody conjugated to Cy3 (diluted 1:250) to label VSV-G-tagged GPCRs expressed on the cell surface. Antibody labeling took place in HEK-complete medium at 37°C and 5% CO_2_ for 30 minutes. Wells were then briefly washed with HEK-complete medium (37°C) and 180µl fresh HEK-complete medium was added. Each well received 20µl of CNO (prepared in HEK-complete medium; 10 times more concentrated than the desired concentration of 10µM), or an equal amount of vehicle (medium only). Cells were then incubated for different time periods (0, 15, 30 or 60 minutes) before fixation. The 0 minute time point was only treated with vehicle and immediately fixed. Vehicle controls were included for each time point to control for the potential contribution of antibody labeling to GPCR internalization. Fixation was carried out with cold 4% (w/v) PFA in PBS for 15 minutes at room temperature. After fixing, cells were washed once with PBS for 3 minutes and subjected to nuclear staining with Hoechst (New England BioLabs; 4082S; diluted 1:5000 in PBS) for 5 minutes. Wells were briefly washed with PBS before adding Antifade mounting medium. Chamber slides were stored at 4°C until imaging. All incubation steps were carried out with 200µl of the respective solution under protection from light to avoid bleaching of Cy3-conjugated antibody and EGFP.

Confocal imaging was performed with a 63x oil immersion objective. For each condition, several images containing one to five cells were acquired with optimal resolution in x, y, and z. Images were processed in Fiji 1.51.f by rolling ball background subtraction followed by a 2x2x2 median filter. Regions-of-interest were generated by cropping individual cells. For analysis, maximum intensity projection of six consecutive z-slices around the center of each cell were obtained. VSV-G and EGFP signals were used to trace the perimeter along the cell surface which separates intra- and extracellular space. A threshold was applied to separate signal from background. The threshold VSV-G area within the perimeter was then measured to quantify internalized GPCRs. To check if this signal is derived from internalized receptors, we confirmed colocalization of VSV-G and EGFP in both channels.

### Analysis of retina transcriptome data

A list of GPCRs was manually collected from Class A (rhodopsin-like, excluding olfactory receptors), Class B (secretin receptor family), Class C (metabotropic glutamate), Class D (fungal mating pheromone receptors), Class E (cAMP receptors), and Class F (Frizzled/Smoothened) and contained in total 361 GPCRs. 58 GPCRs were orphans. Analysis was performed as described in Siegert *et al.* 2012 ^48^.

### Lentiviral vectors

VSV-G enveloped lentiviruses were generated by the Molecular Biology Facility at IST Austria. Briefly, HEK293T cells (5x10^6^) were seeded in 10cm tissue culture dishes and transfected after 24 hours with 6µg packaging plasmid (psPAX2), 2.5µg envelop plasmid (pMD2.G) and 10µg transfer plasmid (PL-SIN-PGK-GPCR-P2A-EGFP-WPRE-miR9T). Culture supernatant containing lentivirus was harvested 24 and 48 hours following transfection. Supernatants from both harvests were pooled, passed through a 0.45µm filter, and stored at -80°C for transduction of HMC3 cells. For primary microglia transduction, supernatants were concentrated through ultracentrifugation (112,000 x g; 1.5h; 4°C) using a 20% sucrose cushion. Pelleted virus was resuspended in PBS and stored at -80°C. For titration of lentivirus preparations, HEK293T cells were seeded into 6-well plates (10^5^ per well) together with a defined volume of virus in various dilutions. After 72 hours, the percentage of EGFP-positive cells was quantified through FACS. Non-transduced cells were used to set the threshold for the EGFP signal. The titer was calculated as transforming units per milliliter (TU/ml) according to the following formula:

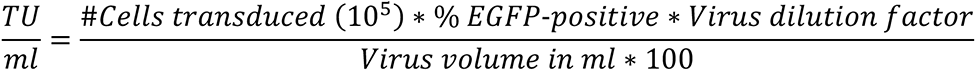

### Generation of HMC3 cell lines stably expressing DREADD-based chimeras

HMC3 cells were seeded into 6-well plates (32,000 per well) together with lentiviral vectors encoding GPCR-P2A-EGFP at a multiplicity of infection (MOI) of 5. Cultures were then expanded for subsequent cell sorting to obtain a pure transduced cell population. For this, cells were trypsinized, pelleted (200 x g; 5min; room temperature) and resuspended in 0.22µm sterile filtered FACS buffer containing 2% (w/v) FBS (Sigma; 12103C; heat-inactivated for 30 minutes at 56°C) and 1mM EDTA in HBSS without Ca^2+^/Mg^2+^. EGFP-positive singlets were sorted into EMEM-complete medium using a Sony SH800SFP cell sorter with a 100µm nozzle chip. Non-transduced cells were used as a negative control to set the threshold for the EGFP signal. The sorting mode was set to “purity” to ensure that only EGFP-expressing cells were included. Culturing of these cells was continued under the previously described maintenance conditions.

### Gene expression profiling in HMC3 cells

Non-transduced HMC3 cells or HMC3 cells stably expressing DREADD-based GPCR chimeras were seeded in 6-well plates at a density of 32,000 cells per well in a total volume of 2ml. Assays were performed three days after seeding when cells were approximately 80% confluent. Cells were then treated by applying fresh EMEM-complete medium containing the respective compounds. Concentrations of levalbuterol, CNO or forskolin were always 10µM. IFNγ/IL1β was added at 10ng/ml each. Untreated control conditions only received fresh EMEM-complete medium. Every experimental repetition included one well per condition. After 6 hours incubation (37°C and 5% CO_2_), wells were briefly washed with DPBS before proceeding with RNA isolation (innuPREP RNA Mini Kit 2.0; Analytik Jena; 845-KS-2040050) according to the manufacturer’s instructions. cDNA was synthesized immediately afterwards (Lunascript RT Super Mix; New England BioLabs; E3010L) with 800-1000ng total RNA as input (same amount for each condition within experimental repetitions) and stored at -20°C.

For gene expression analysis, RT-qPCR (Luna Universal qPCR Master Mix; New England BioLabs; M3003L) was performed in 384 well plates (Bio-Rad; HSR4805) on a Roche Lightcycler 480. Total reaction volume was 10µl containing 1µl of 1:10 diluted cDNA as template and a final concentration of 0.25µM for each primer (**Supplementary Table S3**). Cycle conditions were 60 seconds at 95°C for initial denaturation, followed by 40 cycles of denaturation (15 seconds; 95°C) and annealing/extension (30 seconds; 60°C). Each run was completed with a melting curve analysis to confirm amplification of only one amplicon. Each PCR reaction was run in triplicates from which a mean Cq value was calculated and used for further analysis. dCq values were obtained by normalizing mean Cq values to the geometric mean of four reference genes (GAPDH, ACTB, OAZ1, RPL27) measured within the same sample. ddCq values were then calculated by normalizing dCq values to the respective control condition (untreated cells) within each experimental repetition. Fold changes were obtained by transforming ddCq values from log2-scale to linear scale. These fold changes were used for data visualization after the stimulation with IFNγ/IL1β alone was set to 100% within each experimental repetition.

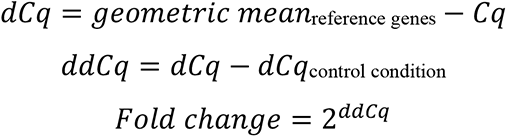

For hierarchical clustering, ratios were calculated between stimulation with IFNγ/IL1β alone, and IFNγ/IL1β together with GPCR ligands (levalbuterol, CNO) or forskolin. Ratios were obtained through dividing linear fold changes by IFNγ/IL1β alone within each experimental repetition. To enable a symmetrical display for up- and downregulation, values of x<1 were converted into negative values by -1/x. These ratios were subsequently subjected to hierarchical clustering in *R* using the *pheatmap* package with default settings.

### Primary microglia cultures

Primary microglia were obtained with adaptations from Bronstein *et al*. ^94^. For one preparation, three to four C57BL6/J mouse pups aged P0-P3 were used. Animals were sprayed with 70% (v/v) ethanol for disinfection before decapitation. Heads were placed in a 10cm on ice containing cold HBSS without Ca^2+^/Mg^2+^. Brains were removed and placed into a fresh 10cm dish with cold HBSS. Under a dissection microscope, meninges were removed before dissecting the cortices, which were subsequently collected in a tube containing 15ml cold HBSS on ice. HBSS was aspirated and 4ml of Trypsin-EDTA (ThermoFisher; 25300-054; 37°C) was added. The tissue was then triturated with a 1000µl pipette tip and incubated at 37°C for 15min in a water bath. Digestion was stopped by adding 4ml of HEK-complete medium (37°C). Samples were pelleted (500 x g for 5min at room temperature) and resuspended in 4ml of HEK-complete medium. The previous centrifugation step was repeated and pellets were resuspended in 10-15ml HEK-complete medium. The cell suspension was passed through a 40µm cell strainer (Szabo Scandic; 352340) and then transferred to a T75 flask to establish a mixed glia culture at 37°C and 5% CO_2_. After three days, medium was replaced with 10ml of fresh HEK-complete medium. Following a total period of 10-14 days after dissection, microglia were harvested from mixed glia cultures through a combination of lidocaine treatment and shaking. A 150mM lidocaine solution (Sigma; L5647) was prepared in HBSS containing Ca^2+^/Mg^2+^ and sterile filtered (0.22µm). Lidocaine was added the T75 flask to a final concentration of 15mM before placing them on a shaker inside a cell culture incubator (37°C and 5% CO_2_) at 70rpm for 25-30min. After this incubation, the supernatant containing detached microglia was collected in a 50ml tube. The flask was briefly washed with 5ml HBSS containing Ca^2+^/Mg^2+^ to gather any remaining microglia and the content was pooled with the previously collected supernatant. EDTA was added to a final concentration of 0.05mM before pelleting microglia (1000 x g; 5min; room temperature). Cells were resuspended in 500µl of HEK-complete medium with a wide 1000µl pipette tip to avoid shear stress. Live cells were counted from a dilution in trypan blue (Sigma; T8154). The concentration was adjusted with HEK-complete medium to 0.2-0.25 million cells/ml to seed approximately 40,000-50,000 cells per well in uncoated 8-well chamber slides (ibidi; 80826; growth area: 1cm^2^) within a total volume of 200µl.

### Live imaging of primary microglia

Primary microglia were transduced with lentiviral vectors encoding for DREADD-β2AR-P2A-EGFP approximately 4-24h after seeding. Virus was applied at an MOI of 0.5-3 which resulted in sparsely transduced cells and live imaging was carried out five to seven days after transduction. Three to four days before live imaging, HEK-complete medium was exchanged with freshly prepared TIC medium optimized for primary microglia culture as described in Bohlen *et al*. ^95^. TIC medium consisted of DMEM/F12 (ThermoFisher; 31331093; with GlutaMAX) containing 5µg/ml N-acetyl-L-cysteine (Sigma; A9165), 5µg/ml bovine insulin (Sigma; I6634), 100µg/ml human apo-transferrin (Sigma; T1147), 100ng/ml sodium selenite (Sigma; S5261), 2ng/ml human TGF-β2 (PepoTech; 100-35B), 100ng/ml murine IL34 (R&D Systems; 5195-ML-010/CF), and 1.5µg/ml ovine wool cholesterol (Sigma; 700000P). For live imaging, primary microglia were labeled with tomato lectin conjugated to DyLight 649 (ThermoFisher; L32472), which was first subjected to ultrafiltration to reduce the concentration of cytotoxic NaN_3_. The required amount of tomato lectin was diluted in PBS (5ml) and applied to a Vivaspin 6 concentrator (Sartorius; VS0601; MWCO: 10kDa; 5ml volume). After centrifugation (4000 x g; 10-15 minutes; 4°C), the concentrate was diluted in DMEM/F12 (37°C) to obtain a final tomato lectin concentration of 5µg/ml (1:200 dilution of stock). Labeling took place for 20min (37°C and 5% CO_2_), after which medium was replaced with 270µl CO_2_-independent Leibovitz’s L15 (ThermoFisher; 21083027; no phenol red; room temperature). Samples were then transferred to a confocal microscope and z-stacks were acquired with a 20x air objective every minute for a total period of 55min at room temperature. In all samples, a tomato lectin and EGFP channel was obtained through simultaneous scanning. The tomato lectin channel was used as autofocus reference which was applied before each z-stack to compensate for vertical drifting. After a 10min baseline recording, a pipette was used to carefully apply 30µl of levalbuterol or CNO (prepared in Leibovitz’s L15; 10 times more concentrated than the desired concentration of 10µM), or an equal amount of vehicle (Leibovitz’s L15 only).

Images were processed in Fiji 1.51 by converting z-stacks to maximum intensity projections, applying a gamma correction of 0.75 for better visualization of faint signals, followed by rolling ball background subtraction and a 1x1 median filter. Regions-of-interest were generated by cropping individual cells. In cases where lateral drifting occurred, image registration was performed with Fiji’s *StackReg* plugin using the *Rigid Body* transformation. The tomato lectin signal was used to quantify changes in cell area for untransduced primary microglia. Microglia transduced with DREADD-β2AR-P2A-EGFP were analyzed through the tomato lectin or EGFP channel, depending on which one provided the best signal. A threshold was applied to separate signal from background. The thresholded area was converted to a binary image and subjected again to a 1x1 median filter to remove unspecific signals. Any remaining signals that did not belong to the respective cell were either removed manually or with Fiji’s *Analyze Particle* function. Subsequently, this binarized area was measured in µm^2^ at all time points during the 55min recording. For the purpose of data visualization, a fold change was calculated for each cell by normalizing area to the mean of the baseline measurement. For final analysis, values from all cells were pooled at representative time points of 1, 5, 10, 25, 40, and 55min.

### Statistical analysis

All analyses were performed with *R*. Data were collected in excel files and imported into R via the *xlsx* or *readxl* package. Linear regression models were generated with the *lme4* package ^96^ and after changing the default contrast for unordered variables (e.g. experimental condition) to “contr.sum”. This allows to run type III Anova on the model to evaluate the overall contribution of unordered effects on the response variable. Post-hoc tests were performed via the *multcomp* package ^97^ with default parameters. If not otherwise indicated, all possible pairwise comparisons were performed. Significance levels are indicated by asterisks (^n.s.^ *p* > 0.05; * *p* ≤ 0.05; ** *p* ≤ 0.01; *** *p* ≤ 0.001). Details about *R* environment, attached packages, statistical models, as well as results of the statistical analysis for each figure are found in **Supplementary Table S4**.

All graphs for data visualization were generated with the *ggplot2* package. Error bars or ribbons represent either standard error of the mean (SEM) calculated by the “mean_se” function (part of *hmisc* package; called through g*gplot2*) or show 95% confidence intervals around a smoothed line generated by the “geom_smooth” function (called through *ggplot2*; using the “loess” method for fitting).

#### Real-time measurement of increases in cAMP levels

We used linear regression to predict the log-transformed luminescence values (baseline-normalized) by an interaction of Time (repeated measurements at regular intervals) and Experimental condition, which is an interaction of Treatment period (Baseline or Ligand), Receptor (GPCR or Empty vector), Ligand (CNO or Levalbuterol) and Concentration (0.01µM, 0.1µM, 1µM or 10µM). A random effect (Experimental repetition) was included to account for the dependency of data, which results from repeated measurements within each individual experiment. This model was used to test whether ligand treatment of individual Receptor-Ligand-Concentration interactions results in significant differences from the baseline measurement.

#### SRE reporter assay

We used a two-sided one-sample T-test to investigate whether ligand-treated conditions are significantly different from a value of 1, which represents the vehicle control of the respective experimental repetition.

#### Real-time assay measurement of β-arrestin 2 recruitment

We used linear regression to predict luminescence values (normalized to baseline and further to the respective vehicle control) by and interaction of Time (repeated measurements at regular intervals) and Experimental condition, which is an interaction of Treatment period (Baseline or Ligand) and Receptor (β2AR-SmBiT or DREADD-β2AR-SmBiT). A random effect (Experimental repetition) was included to account for the dependency of data, which results from repeated measurements within each individual experiment. This model was used to test whether ligand treatment of each Receptor results in significant differences from the baseline measurement.

#### Western blot to quantify ERK1/2 phosphorylation

We used linear regression to predict ratios between pERK1/2 and GAPDH by Experimental condition, which represents different treatment durations (untreated, 2min, 5min, or 15min). A random effect (Experimental repetition) was included to account for the dependency of data that are derived from the same experimental repetition.

#### GPCR internalization assay

We used linear regression to predict internalized area in µm^2^ (measured individually per cell based on thresholded VSV-G signal) by Experimental condition, which is an interaction of Ligand (CNO or Vehicle) and Treatment period (0min, 15min, 30min or 60min). This model was used to test whether CNO treatment shows significant differences from vehicle controls at corresponding Treatment periods.

#### Real-time measurement of decreases in cAMP levels

We used linear regression to predict the log-transformed luminescence values (normalized to baseline and further to the respective vehicle control) by Experimental condition, which is an interaction of Treatment period (Baseline, Ligand or Forskolin) and Receptor (GPCR or Empty vector). A random effect (Experimental repetition) was included to account for the dependency of data, which results from repeated measurements within each individual experiment. This model was used to test Receptor and Empty vector for significant differences between their three Treatment periods. The time intervals of repeated measurements were not included as a predictor in the model as it was not necessary to improve the fit. This is because measured values are rather uniformly distributed within the three different Treatment periods, meaning that this variably can already explain most of the variability in the data.

#### CRE reporter assay

We used a two-sided one-sample T-test to investigate whether ligand-treated conditions are significantly different from a value of 1, which represents the vehicle control of the respective experimental repetition.

#### Gene expression profiling in HMC3 cells

We used a two-sided one-sample T-test to confirm that recombinant cytokine stimulation induces inflammatory gene expression. We compared the linear fold change of stimulated conditions to a value of 1, which represents the untreated control of the respective experimental repetition. For further comparison of different treatment conditions, we used linear regression. We predicted ddCq values for individual transcripts by Experimental condition, which represents the treatment with different compounds alone or in combination. A random effect (Experimental repetition) was included to account for the dependency of data derived from the same experimental repetition. Separate models were generated for each investigated transcript (IL6, TNF and IL1β) and cell line (non-transduced, DREADD-β2AR, DREADD-GPR65, and DREADD-GPR109A). These models were used to test for significant differences between different treatments.

#### Live imaging of primary microglia

We used linear regression to model the change of total cell area in µm^2^ by using an interaction of the two predictors Time and Experimental condition. Experimental condition itself is an interaction of Treatment period (Baseline or Ligand) and Experimental group (Untransduced_Levalbuterol, DREADD-β2AR_CNO, Untransduced_Vehicle, or Untransduced_CNO). A random effect (Cell ID) was included to account for the dependency of data, which results from repeated measurements on individual cells. This random effect also accounts for size differences between cells. This model was used to test for significant differences between the two Treatment periods within each Experimental group.

---

**Supplementary Figure S1:**
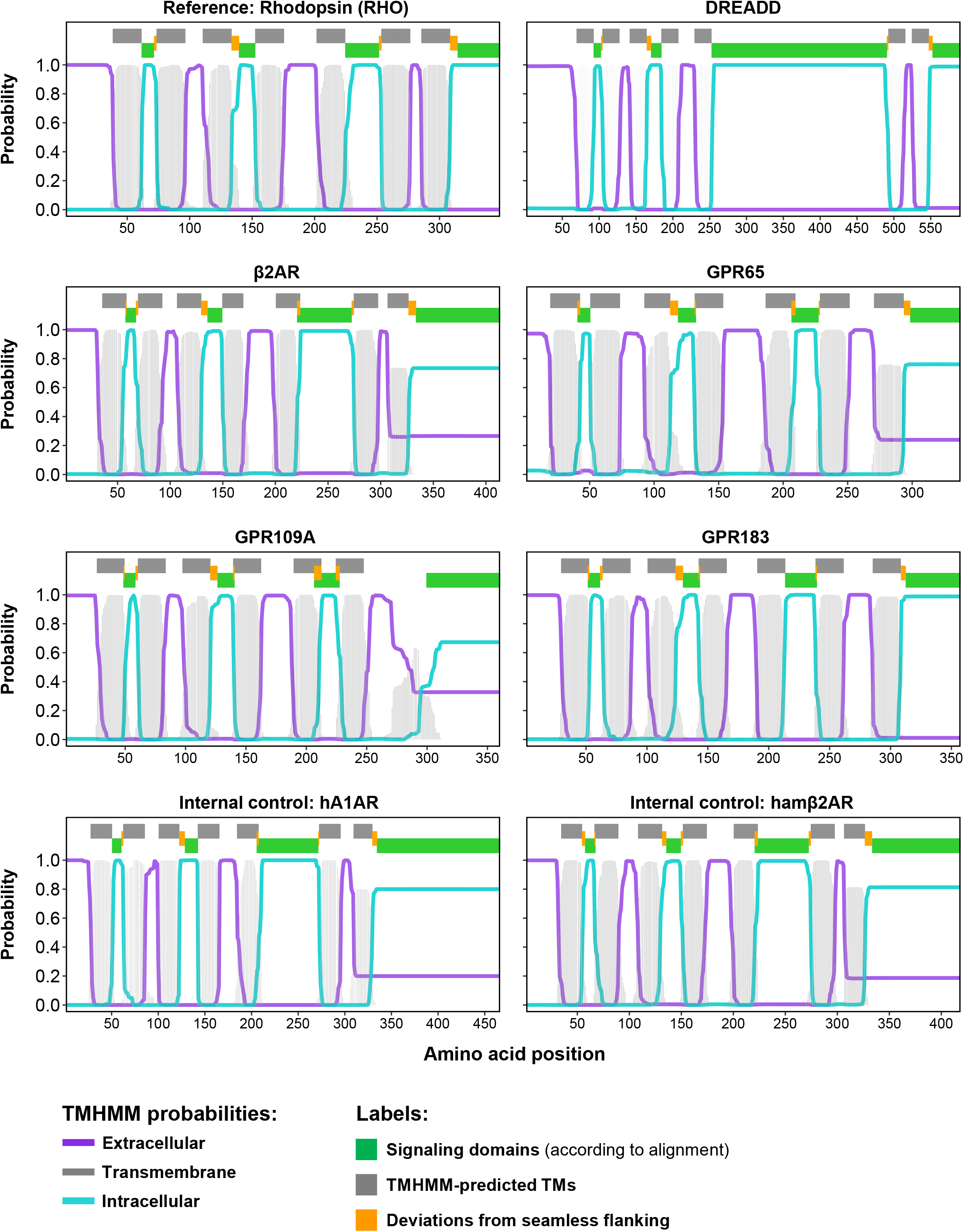
TMHMM-predictions support multiple protein sequence alignment accuracy. TMHMM probability plots for rhodopsin (RHO), DREADD, four representative GPCRs-of-interest, human α1-adrenergic receptor (hα1AR), and hamster β2-adrenergic receptor (hamβ2AR). TMHMM reports the probability that a GPCR sequence is located in the extracellular (purple line), transmembrane (light grey line), or intracellular (light turquoise line) space. The top of each plot shows the results from multiple protein sequence alignment from N-to C-terminus. Alignment-identified intracellular signaling domains (green bars) were tightly flanked by TMHMM-predicted transmembrane domains (TMs; dark grey bars). Deviations from seamless flanking (orange bars) were minimal and comparable to deviations within the alignment reference (rhodopsin) and internal controls (hα1AR, hamβ2AR).

**Supplementary Figure S2:**
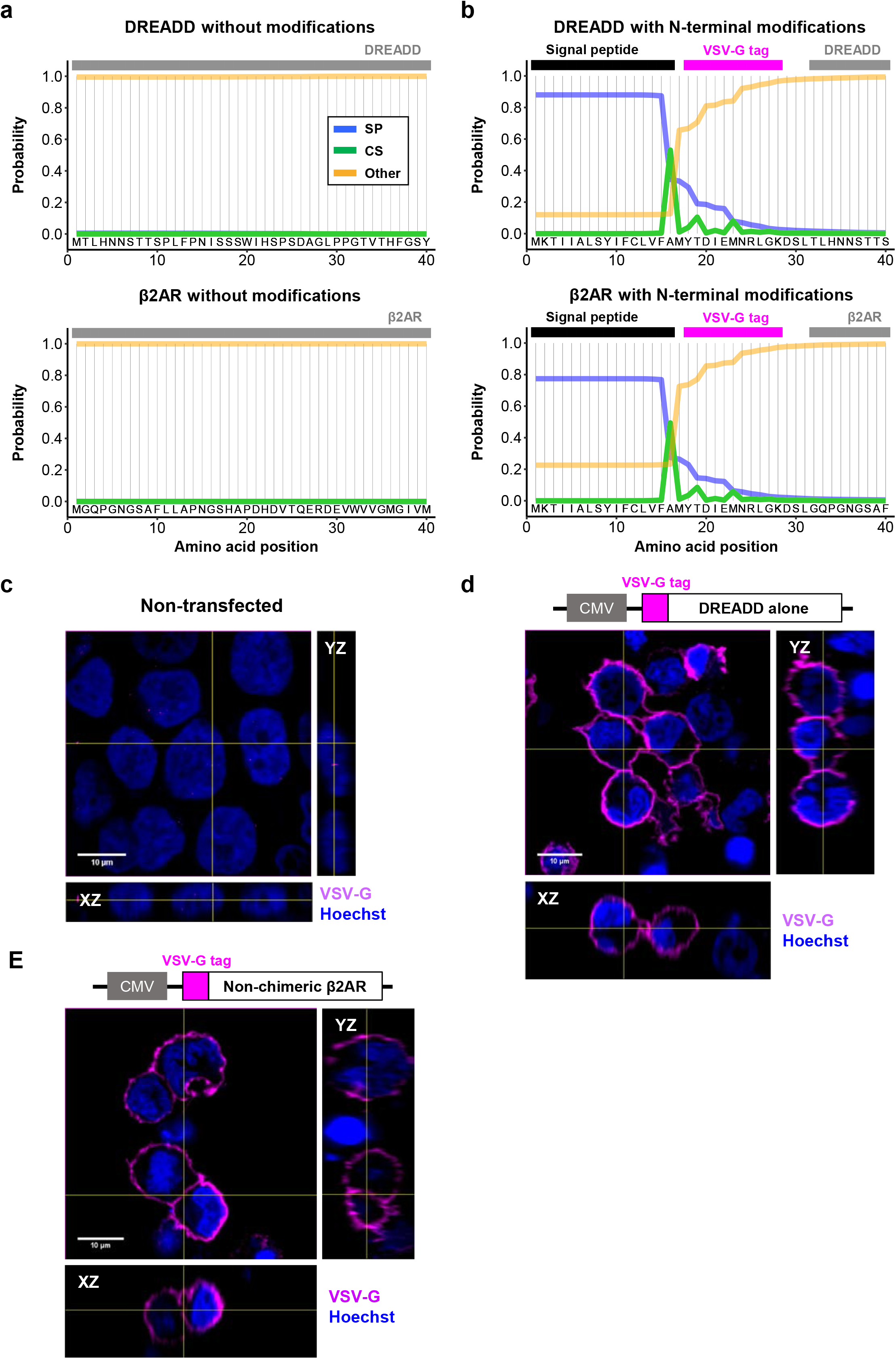
N-terminal modifications of DREADD and β2AR are compatible with cell surface expression. a-b: SignalP probability plots for DREADD (top) and β2AR (below) without any modification (**a**) or with signal peptide and VSV-G epitope tag (**b**). Blue line (SP): probability that sequence belongs to a signal peptide. Green line (CS): probability of signal peptide cleavage. Orange line (Other): probability that sequence does not contain a signal peptide. Grey bar represents protein sequence of either DREADD or β2AR. **c-e:** Orthogonal views of HEK cells immunostained for VSV-G tag under non-permeabilizing conditions in non-transfected cells (**c**), and cells transfected with DREADD alone (**d**) or non-chimeric β2AR (**e**) transfected cells. Magenta: VSV-G tag. Blue: nuclear staining with Hoechst. CMV, human cytomegalovirus promoter.

**Supplementary Figure S3:**
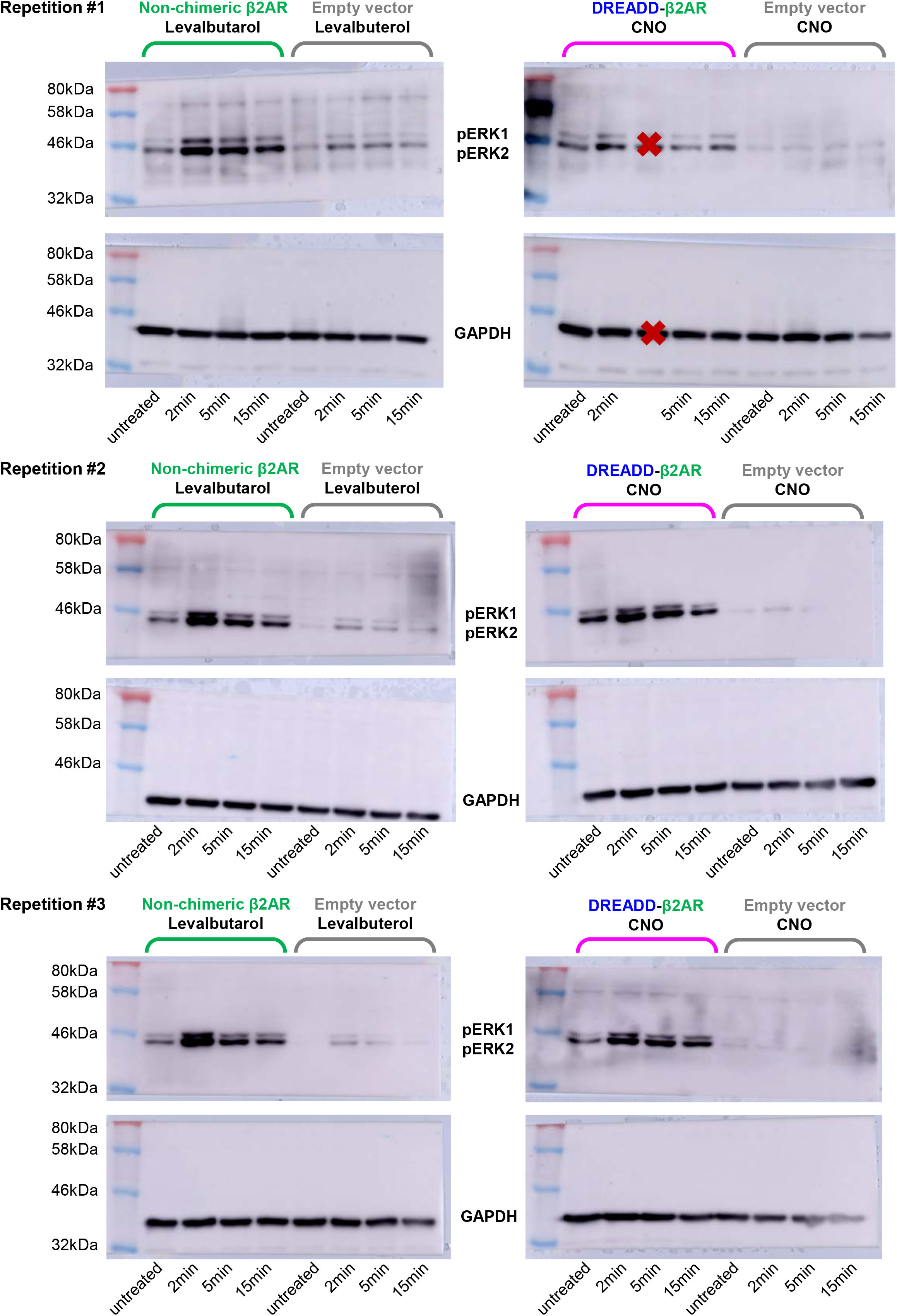
Western blot analysis of ERK1/2 phosphorylation. Summary of all Western blots for phosphorylation analysis of extracellular signal-regulated kinases 1 and 2 (ERK1/2) in untreated, levalbuterol- or CNO-treated HEK cells transfected with non-chimeric β2AR (green), DREADD-β2AR (magenta), or empty vector (grey). Densitometry analysis shown in **Figure 3C**. Red cross: lanes excluded due to incorrect sample loading.

**Supplementary Figure S4:**
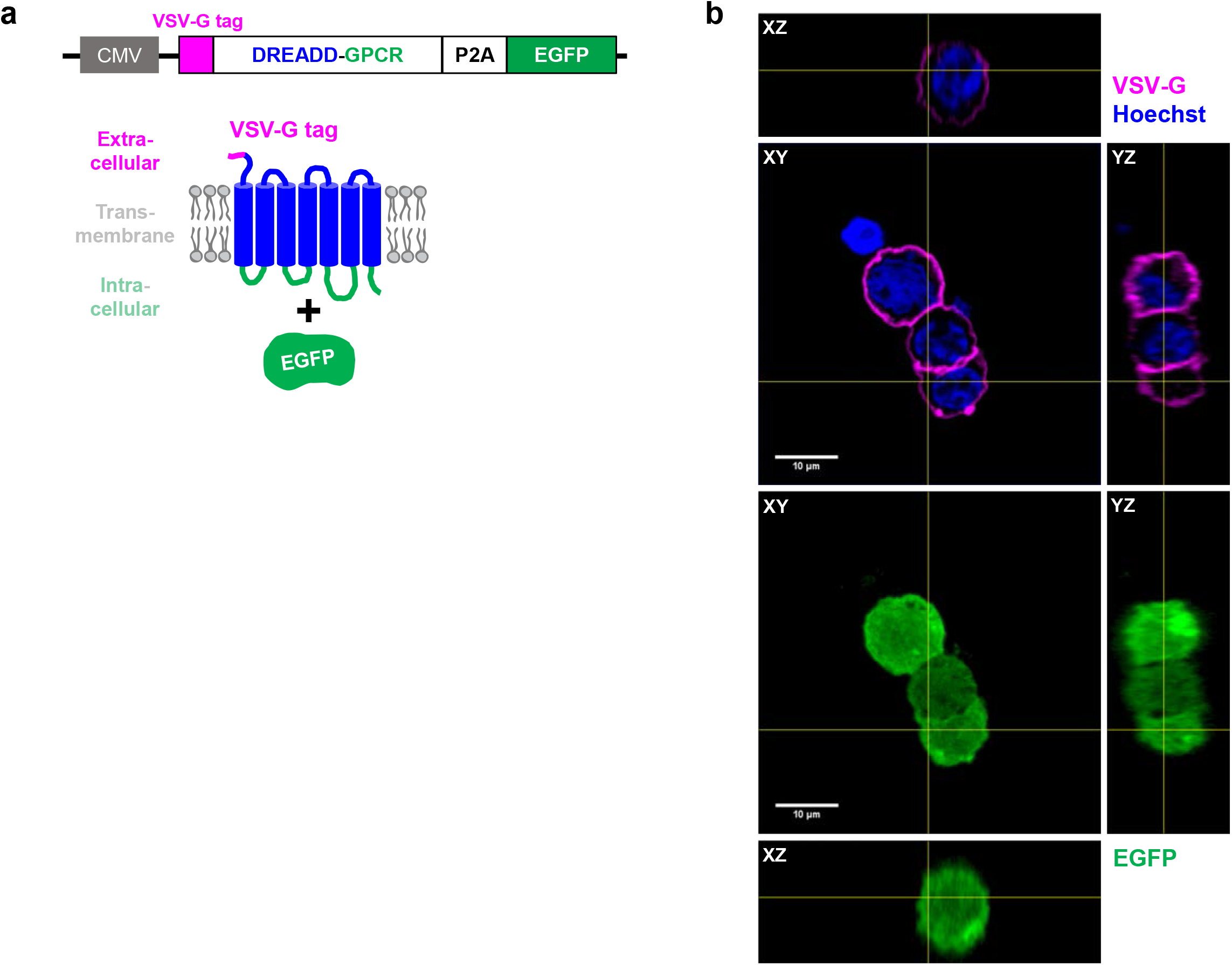
Vector validation for bicistronic expression of DREADD-GPCRs and EGFP. **a:** Schematic of a bicistronic vector with human cytomegalovirus (CMV) promoter, VSV-G epitope, DREADD-GPCR, the self-cleaving P2A peptide sequence, and EGFP. Transfected cells express the GPCR on the cell surface and EGFP in the cytoplasm. **b:** Orthogonal view of HEK cells transfected with DREADD-β2AR-P2A-EGFP immunostained for the VSV-G tag under non-permeabilizing conditions. Magenta: VSV-G tag. Green: EGFP. Blue: nuclear staining with Hoechst.

**Supplementary Figure S5:**
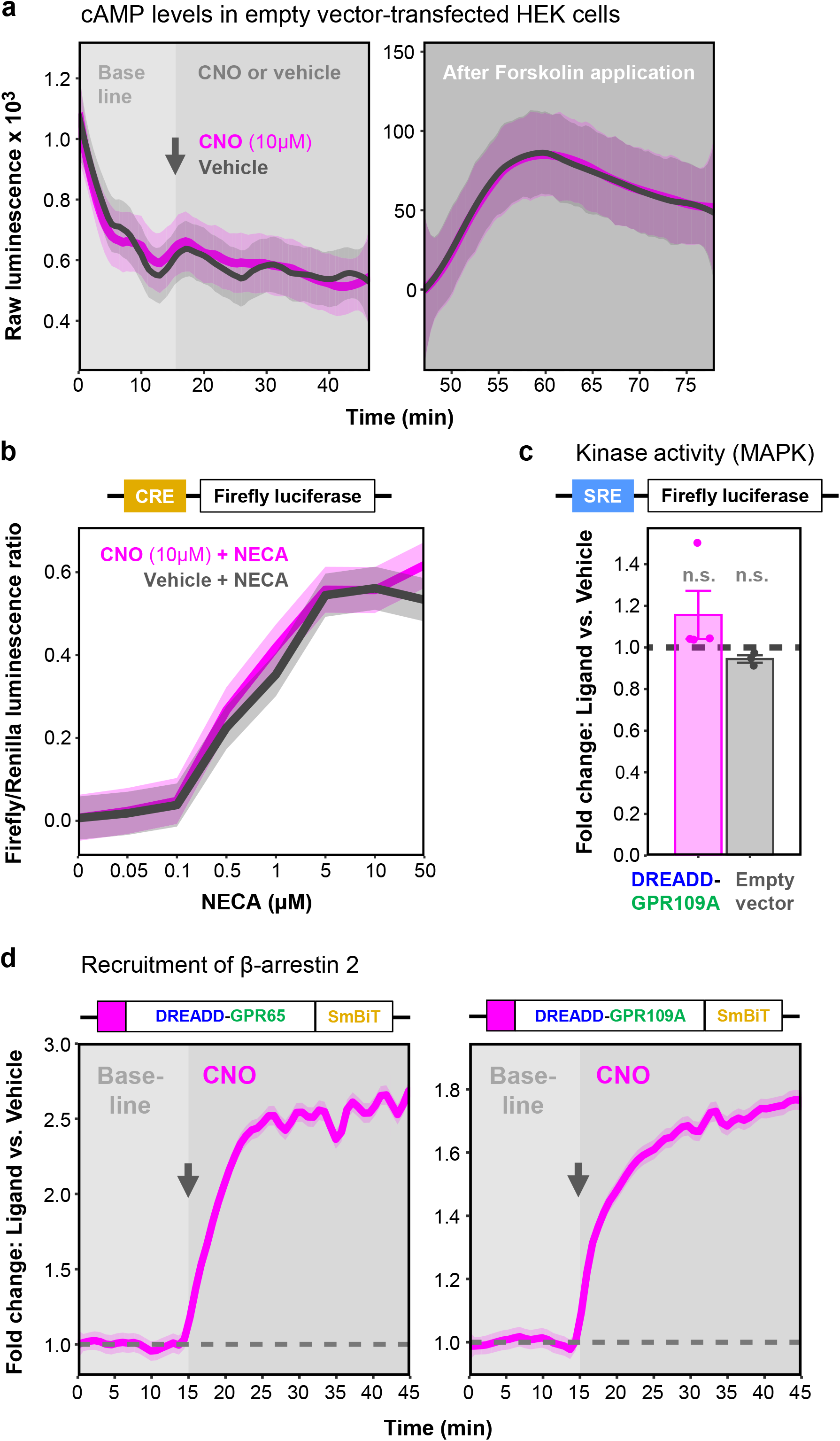
Additional controls and validations. **a:** CNO does not impair forskolin-induced cAMP synthesis. Real-time measurement of cAMP-dependent luciferase activity in HEK cells transfected with empty vector. Baseline measurements followed by application of CNO or vehicle (grey arrow for onset) and forskolin (second panel). Graph shows raw luminescence units of cells treated with CNO (magenta) or vehicle (grey). Ribbons: 95% confidence intervals of five repetitions. **b:** CNO does not impact NECA-induced transcription from a cAMP responsive element (CRE). Endpoint measurement of cAMP responsive element (CRE)-dependent firefly luciferase activity in HEK cells transfected with empty vector. Treatment with increasing concentrations of NECA in presence of either CNO (magenta) or vehicle (grey). Graph shows firefly/renilla luciferase luminescence ratios. Ribbons: standard error of the mean of three technical replicates from one representative experiment. **c:** DREADD-GPR109A does not influence SRE reporter activity. Endpoint measurement of serum responsive element (SRE)-dependent luciferase activity in HEK cells transfected with DREADD-GPR109A (magenta) or empty vector (grey). Ligand stimulation with CNO. Dashed line: level of the respective vehicle control. Error bars: standard error of the mean of three to four repetitions. One-sample T-test: *p^ns^* > 0.05. **d:** DREADD-GPR65 and GPR109A recruit β-arrestin 2. Real-time measurement of β-arrestin 2 recruitment in HEK cells transfected with DREADD-GPR65 (left) or DREADD-GPR109A (right). Baseline measurements followed by CNO application (grey arrow shows onset). Graphs show baseline-normalized fold changes compared to vehicle. Dashed line: level of the vehicle control. Ribbons: 95% confidence intervals of three technical replicates from one representative experiment.

**Supplementary Figure S6:**
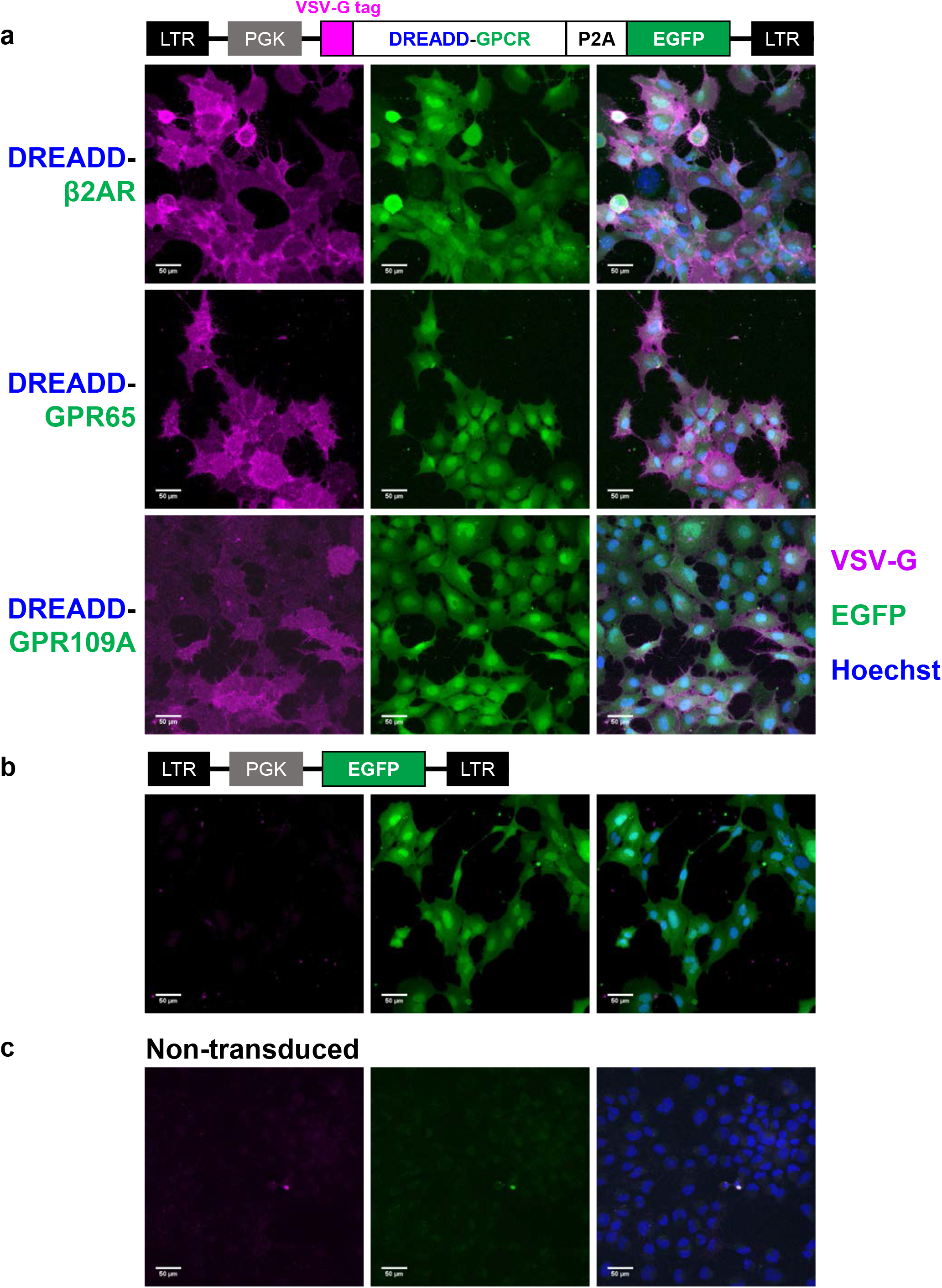
Stable expression of DREADD-based chimeras in HMC3 cells through lentiviral vectors. **a:** Schematic of lentiviral vector with DREADD-GPCR-P2A-EGFP flanked by long terminal repeats (LTRs) and expression driven by the phosphoglycerate kinase (PGK) promoter. Images show maximum intensity projections of HMC3 cells immunostained for the VSV-G tag under non-permeabilizing conditions. Cells were either transduced with DREADD-β2AR, DREADD-GPR65, or DREADD-GPR109A (**a**); a control vector encoding only EGFP (**b**); or remained non-transduced (**c**). Magenta: VSV-G. Green: EGFP. Blue: nuclear staining with Hoechst.

**Supplementary Figure S7:**
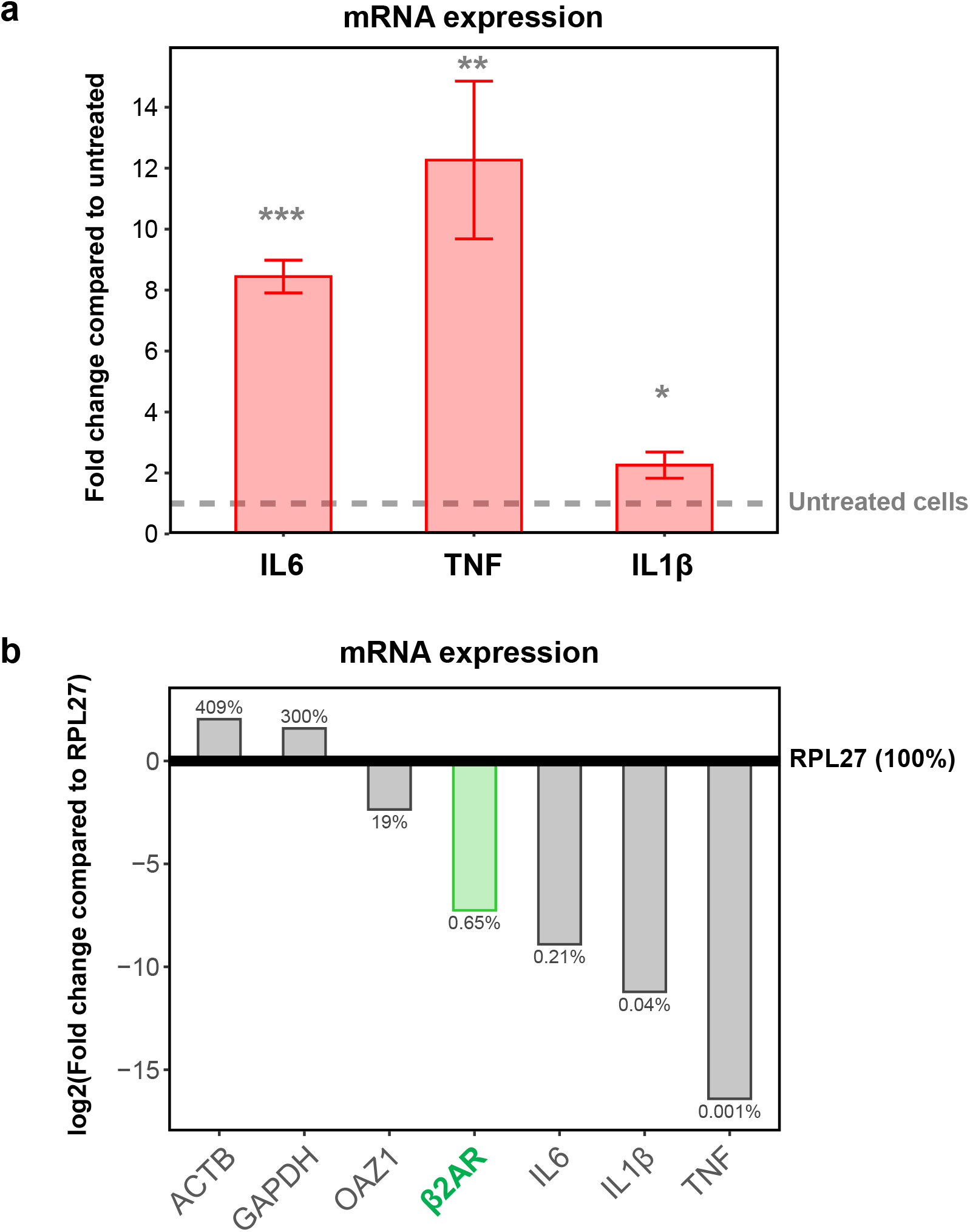
HMC3 cells respond to the recombinant cytokines IFNγ and IL1β and endogenously express β2AR. **a:** Quantitative reverse transcription PCR (RT-qPCR) of HMC3 cells exposed to recombinant interferon γ (IFNγ) and interleukin 1β (IL1β). Expression values of interleukin 6 (IL6), tumor necrosis factor (TNF), and IL1β shown as fold change compared to untreated cells (dashed line). Error bars: standard error of the mean of eight repetitions. One-sample T-test: *p\*\*\** < 0.001; *p\** < 0.01; *p\** < 0.05. **b:** RT-qPCR-based comparison of gene expression. Bars: log2-fold changes compared to RPL27 and are based on the mean of three technical replicates from a representative experiment. Relative transcript abundance is additionally shown as a percentage of RPL27 (100%). ACTB, β-actin. GAPDH, glyceraldehyde-3-phosphate dehydrogenase. OAZ1, ornithine decarboxylase antizyme 1. RPL27, ribosomal protein L27.

**Supplementary Figure S8:**
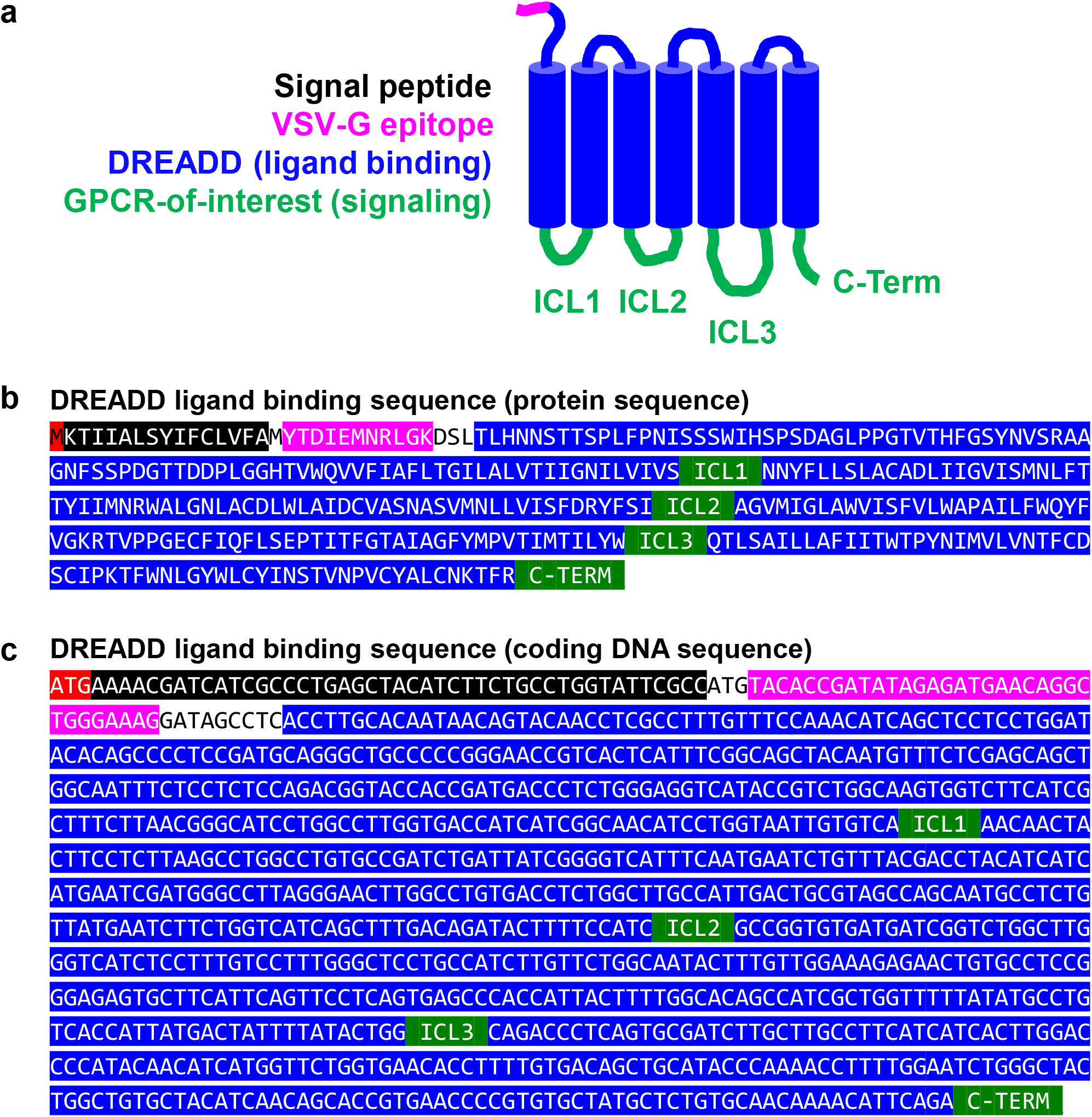
*In-silico* design of DREADD-based chimeras for a GPCR-of-interest. **a:** Schematic of a DREADD-based GPCR chimera containing signal peptide (black) and VSV-G epitope tag (magenta) at the N-terminus. Blue: ligand binding domains of DREADD. Green: signaling domains of a GPCR-of-interest. **b-c:** Protein (**b**) and corresponding coding DNA sequence (**c**) of the DREADD ligand binding domains. To generate a DREADD-based chimera for a specific GPCR-of-interest, sequences encoding for the respective signaling domains (green) are inserted in the designated fields. ICL1-3, intracellular loops 1-3. C-Term, C-terminus. Protein sequences of putative signaling domains of 292 GPCRs-of-interest are available in **Supplementary Table S1**.

**Supplementary Table S1.** Library of putative signaling domains of GPCRs-of-interest. Species indicators: b, bovine; ham, hamster; h, human; m, mouse; r, rat. All GPCRs without a species indicator are human. GPCRs with *: DREADDs.

**Supplementary Table S2.** HEK cell transfection scheme.

**Supplementary Table S3.** List of RT-qPCR primers.

**Supplementary Table S4.** Summary of statistical analysis.

